# Oncomodulin Regulates Spontaneous Calcium Signaling and Maturation of Afferent Innervation in Cochlear Outer Hair Cells

**DOI:** 10.1101/2023.03.01.529895

**Authors:** Yang Yang, Kaitlin Murtha, Leslie K. Climer, Federico Ceriani, Pierce Thompson, Aubrey J. Hornak, Walter Marcotti, Dwayne D. Simmons

**Affiliations:** Department of Biology, Baylor University, 101 Bagby Ave, Waco, TX; School of Biosciences, University of Sheffield, S10 2TN Sheffield, United Kingdom; Department of Integrative Biology and Physiology, University of California, Los Angeles, CA; Sheffield Neuroscience Institute, University of Sheffield, Sheffield, S10 2TN, UK; Department of Psychology and Neuroscience, Baylor University, Waco, TX

**Keywords:** Oncomodulin (OCM), Outer Hair Cell (OHC), Purinergic receptors, Ribbon synapse, Spontaneous Calcium activity

## Abstract

Cochlear outer hair cells (OHCs) are responsible for the exquisite frequency selectivity and sensitivity of mammalian hearing. During development, the maturation of OHC afferent connectivity is refined by coordinated spontaneous Ca^2+^ activity in both sensory and non-sensory cells. Calcium signaling in neonatal OHCs can be modulated by Oncomodulin (OCM, β-parvalbumin), an EF-hand calcium-binding protein. Here, we investigated whether OCM regulates OHC spontaneous Ca^2+^ activity and afferent connectivity during development. Using a genetically encoded Ca^2+^ sensor (GCaMP6s) expressed in OHCs in wild-type (Ocm^+/+^) and Ocm knockout (Ocm^-/-^) littermates, we found increased spontaneous Ca^2+^ activity and upregulation of purinergic receptors in OHCs from GCaMP6s Ocm^-/-^ cochlea immediately following birth. The afferent synaptic maturation of OHCs was delayed in the absence of OCM, leading to an increased number of ribbon synapses and afferent fibers on GCaMP6s Ocm^-/-^ OHCs before hearing onset. We propose that OCM regulates the spontaneous Ca^2+^ signaling in the developing cochlea and the maturation of OHC afferent innervation.

## Introduction

The regulation and control of Ca^2+^ is a major challenge for cochlear sensory cells during development as well as in the adult. Similar to other sensory systems, the developing cochlea exhibits intrinsically generated, sound-independent spontaneous Ca^2+^ activity that is critical for the maturation and refinement of neural circuits (Babola et al., 2021; Blankenship and Feller, 2010; Clause et al., 2014; Lippe, 1994). The mammalian cochlea has two types of specialized sensory cells, which are involved in the transduction of sound into electrical responses (Dallos, 1992). Inner hair cells (IHCs) relay sound information via glutaminergic synapses onto type I afferent spiral ganglion neurons. Outer hair cells (OHCs) enhance cochlear sensitivity and frequency tuning of the cochlear partition and are primarily innervated by cholinergic medial olivocochlear neurons that form a sound-evoked acoustic reflex (Guinan, 2018). Additionally, OHCs form synapses onto type II afferent spiral ganglion neurons that may be activated by traumatic noise exposures (Flores et al., 2015; Liu et al., 2015). In rodents, the maturation of OHC innervation patterns occurs during the first and second postnatal weeks (Simmons, 1994; Simmons et al., 1996). Recent studies show that the OHC innervation is refined by coordinated spontaneous Ca^2+^ activity through the modulation of voltage-gated Ca^2+^ channels and purinergic receptor signaling (Ceriani et al., 2019; Jeng et al., 2020). However, little is known about the intracellular Ca^2+^ signaling network that modulates OHC spontaneous Ca^2+^ activity during cochlear maturation.

A multitude of transporters, pumps, exchangers, and calcium-binding proteins (CaBPs) are integral to the Ca^2+^ signaling network in OHCs and tightly regulate Ca^2+^ activity and homeostasis. Oncomodulin (OCM), a small EF-hand CaBP of approximately 12 kDa, is the β isoform of parvalbumin and shares at least 53% sequence identity with α-parvalbumin (PVALB) (Banville and Boie, 1989). Previous studies show that in the cochlea, OCM is expressed specifically by OHCs and preferentially localizes to the lateral membrane, the basal portion of the hair bundle, and the basal pole adjacent to efferent terminals (Simmons et al., 2010). After hearing onset, which in most altricial rodents occurs at around postnatal day 12 (P12), OHCs express high levels of OCM (2-3 mM) compared to other CaBPs (Hackney et al., 2005). Together, this suggests that OCM may have a unique function relative to the other CaBPs in OHCs. OCM is an important CaBP for which targeted deletion causes hearing loss (Pangrsic et al., 2015; Tong et al., 2016). The absence of OCM uniquely alters Ca^2+^ signaling in OHCs before the onset of hearing (Murtha et al., 2022). Given that spontaneous Ca^2+^ activity plays a major role in OHC development, we hypothesized that OCM modulates spontaneous Ca^2+^ activity during development.

To investigate how OCM regulates spontaneous Ca^2+^ activity in developing OHCs, we expressed a genetically encoded, tissue-specific Ca^2+^ sensor (*Atoh1*-GCaMP6s) in *Ocm* wild-type (*Ocm*^*+/+*^) and *Ocm* knockout (*Ocm*^*-/-*^) mice. GCaMP6s *Ocm*^*-/-*^ mice exhibited early onset hearing loss, and their OHCs showed faster KCl-induced Ca^2+^ transients than those recorded from GCaMP6s *Ocm*^*+/+*^ mice and other mouse strains (Climer et al., 2021; Murtha et al., 2022; Tong et al., 2016). In neonatal mice (P2), we observed spontaneous Ca^2+^ activity in OHCs that was synchronized by Ca^2+^ waves elicited in the greater epithelial ridge (GER). However, GCaMP6s *Ocm*^*-/-*^ OHCs exhibited higher synchronization and stronger fractional change of GCaMP6s fluorescence intensity (∆F/F_0_) during Ca^2+^ activity in the GER, compared to GCaMP6s *Ocm*^*+/+*^ OHCs. We also found that the expression of P2X2, one of the main purinergic receptors in the cochlea, was upregulated in GCaMP6s *Ocm*^*-/-*^ cochlea. GCaMP6s *Ocm*^*-/-*^ OHCs showed delayed synaptic pruning with an increased number of tunnel crossing fibers during development. We propose that the lack of OCM alters the spontaneous Ca^2+^ activity via ATP signaling and affects the afferent maturation and innervation of the OHCs.

## Results

### Lack of OCM expression causes early hearing loss in GCaMP6s adult mice

We utilized tissue-specific expression of GCaMP6s (**Figure 1A**) to investigate Ca^2+^ signaling in the cochlea. *Atoh1*-driven Cre mice were crossed with the Ai96 mice containing a floxed-STOP cassette GCaMP6s. *Atoh1* expression is found in IHCs and OHCs during development (Mulvaney and Dabdoub, 2012). As a sensitive GFP-based Ca^2+^ sensor, GCaMP6s has been used to probe fast Ca^2+^ dynamics and low peak Ca^2+^ accumulations in neurons (Chen et al., 2013; Lukasz and Kindt, 2018; Shilling-Scrivo et al., 2021). Calcium signaling in the OHCs was investigated using GCaMP6s *Ocm* wildtype (*Ocm*^*+/+*^) and *Ocm* knockout (*Ocm*^*-/-*^) mice (**Figure 1A-B**, see Materials and Methods). Initially, we examined OHC function in adult *Ocm* mice expressing GCaMP6s by measuring distortion product otoacoustic emissions (DPOAEs). At 3-4 weeks (wks), there were no differences in DPOAE thresholds between GCaMP6s *Ocm*^*+/+*^ and GCaMP6s *Ocm*^*-/-*^ mice (*P* > 0.90, *two-way ANOVA*, **Figure 1C**). However, by 7-9 wks, GCaMP6s *Ocm*^*-/-*^ mice showed hearing loss with higher DPOAE thresholds at 16, 22, and 32 kHz (*P* < 0.05, *two-way ANOVA*, **Figure 1D**). These results are consistent with other studies showing that OCM is critical for maintaining cochlear function in adult mice (Climer et al., 2021; Tong et al., 2016).

**Figure 1.**
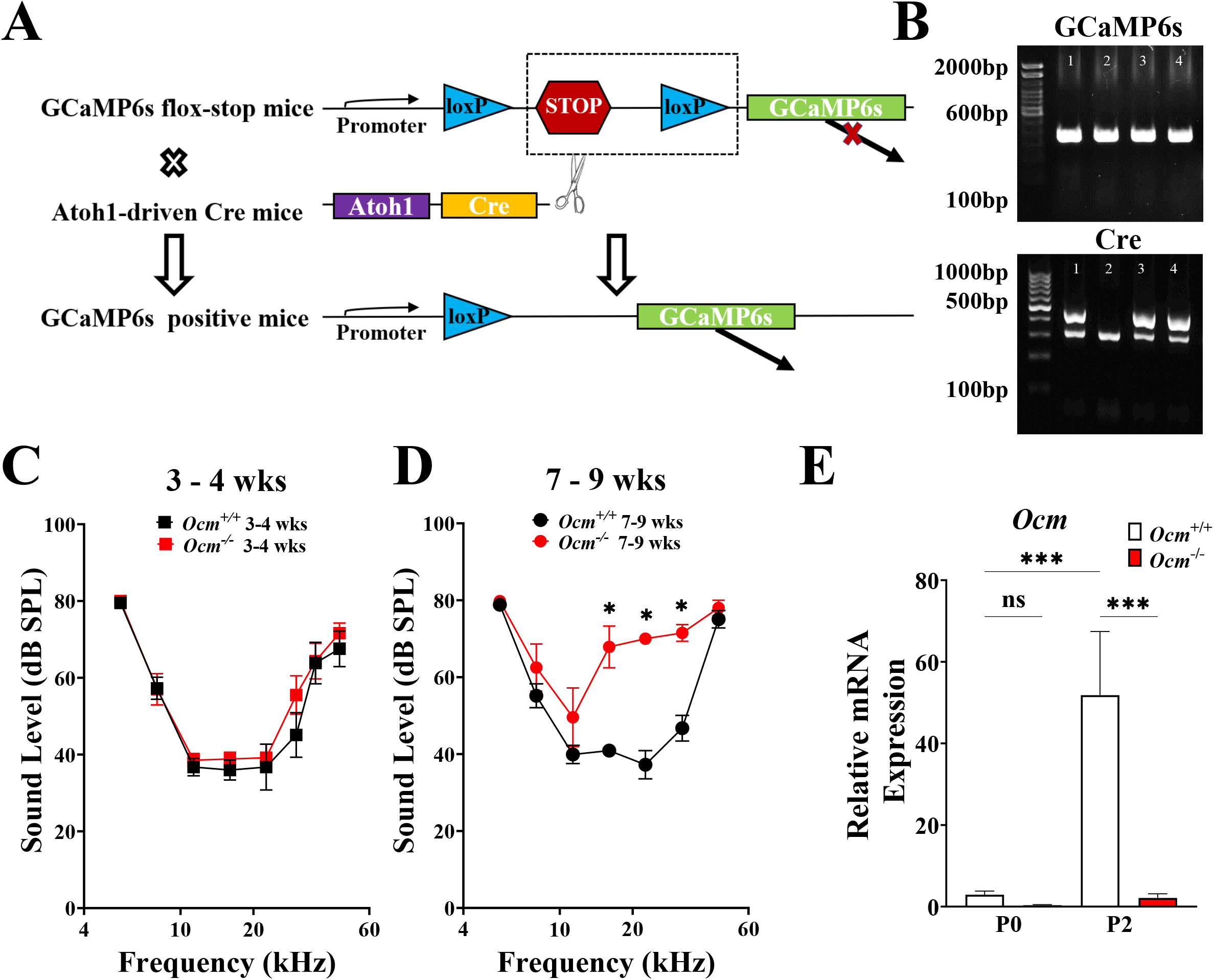
Genetic deletion of *Ocm* or stable expression of GCaMP6s in OHCs showed no phenotypic difference in hearing at 3-4 weeks of age. **A**. Schematic showing generation of *Atoh1*-driven, GCaMP6s *Ocm*^*+/+*,^ and *Ocm*^*-/-*^ mice. Ai96(RCL-GCaMP6s, The Jackson Laboratory) mice contain a floxed-STOP cassette preventing transcription of the GCaMP6. After crossing with *Atoh1*-driven Cre mice, the floxed-STOP cassette was removed, leading to a tissue-specific expression of GCaMP6s in the inner ear region. **B**. Confirmation of Cre-driven, genetically encoded GCaMP6s. **C-D**. Distortion product otoacoustic emissions (DPOAEs) were measured from 3-4 weeks (wks) and 7-9 wks GCaMP6s *Ocm*^*-/-*^ mice and aged-matched GCaMP6s *Ocm*^*+/+*^ mice (6 *Ocm*^*+/+*^ mice, 8 *Ocm*^*-/-*^ mice at 3-4 wks, 11 *Ocm*^*+/+*^ mice, 5 *Ocm*^*-/-*^ mice at 7-9 wks were tested). *: *P* < 0.05, *two-way ANOVA*, P = 0.63 at 5kHz, P = 0.98 at 8kHz, P = 0.89 at 11kHz, P = 0.99 at 45kHz, P < 0.05 at 16, 22 and 32kHZ, Bonferroni’s post-test. **E**. qRT-PCR results of *Ocm* mRNA expression from GCaMP6s *Ocm*^*+/+*^ and *Ocm*^*-/-*^ cochlea at postnatal day 0 (P0) and P2 (n ≥ 3 for each genotype at each age). Results are mean (± SEM) and normalized to *Ocm*^*+/+*^at P0, *: *P* < 0.05, one-way ANOVA, P > 0.99 for P0 GCaMP6s *Ocm*^+/+^ vs. P0 GCaMP6s *Ocm*^-/-^, P < 0.001 for P2 GCaMP6s *Ocm*^+/+^ vs. P2 GCaMP6s *Ocm*^-/-^ and P0 GCaMP6s *Ocm*^+/+^ vs. P2 GCaMP6s *Ocm*^+/+^, Bonferroni’s post-test.

In the cochlea of GCaMP6s *Ocm*^+/+^ mice, *Ocm* mRNA was detected as early as P0 and was significantly upregulated at P2. Relative *Ocm* mRNA expression in GCaMP6s *Ocm*^*-/-*^ cochlea was negligible compared to that of GCaMP6s *Ocm*^+/+^ (**Figure 1E**, *P* < 0.001, *one-way ANOVA*). Using confocal microscopy, we found that at P2, GCaMP6s *Ocm*^*+/+*,^ and GCaMP6s *Ocm*^*-/-*^ mice showed endogenous GCaMP6s fluorescence in the cochlea (**Figure 1F**). GCaMP6s *Ocm*^*-/-*^ OHCs exhibited higher baseline fluorescence intensity compared to GCaMP6s *Ocm*^+/+^ OHCs (F_0_, **Figure 2**), indicating a possible higher basal level of intracellular Ca^2+^ due to the lack of OCM. A similar result has been reported using the ratiometric dye Fura 2 (Murtha et al., 2022). OCM can be detected in P2 GCaMP6s *Ocm*^+/+^ OHCs (**Figure 2**). Moreover, we found a gradient expression of OCM along the tonotopic axis of the cochlea of P2 GCaMP6s mice, with a higher OCM level in high-frequency OHC (base, **Figure 2**). These results confirm that OCM protein can be detected as early as P2 in OHCs, similar to other mouse strains (Climer et al., 2019; Murtha et al., 2022).

**Figure 2.**
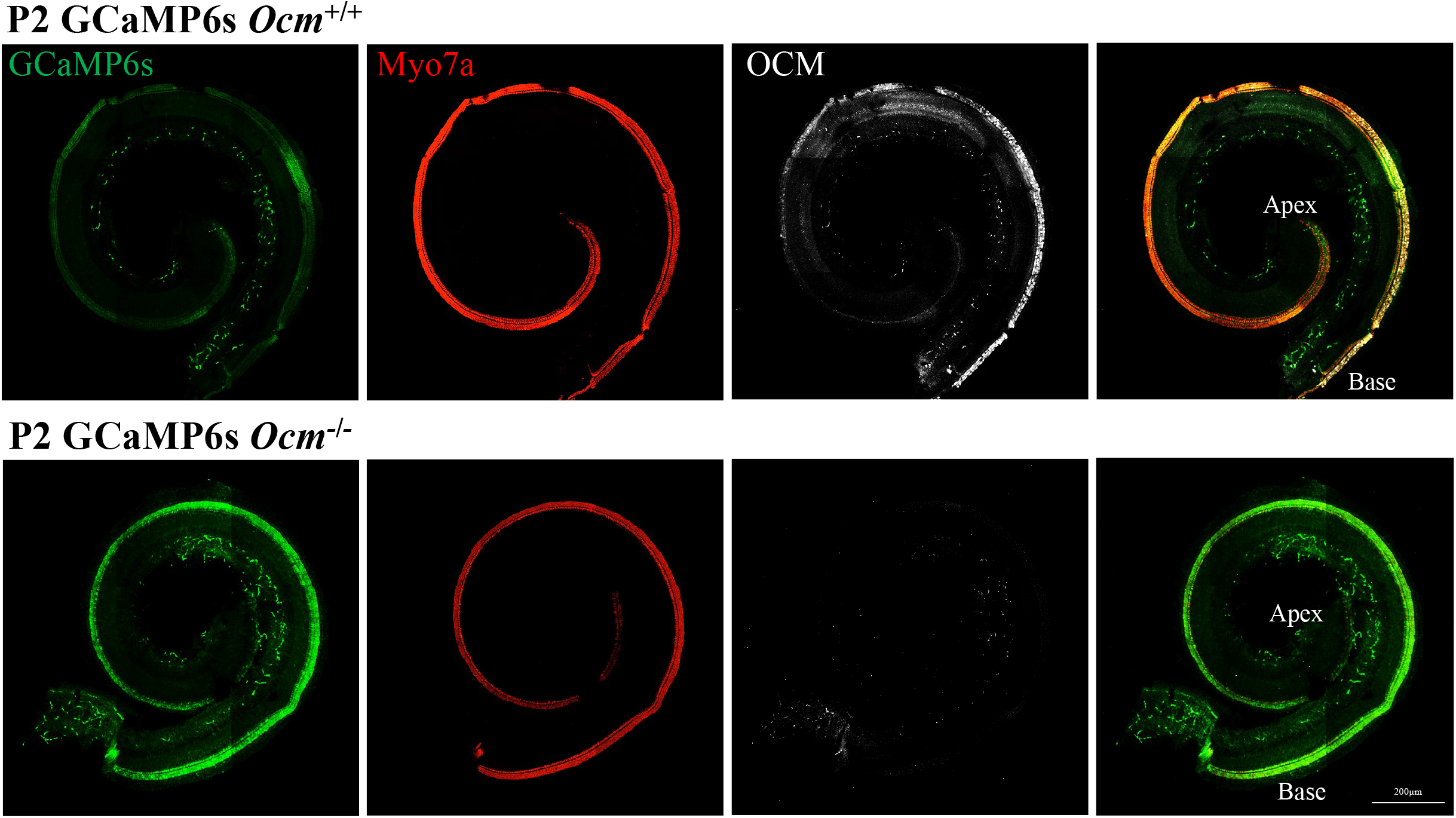
Gradient expression of OCM along the tonotopic axis in P2 GCaMP6s mice. Maximum intensity projections of confocal 9 tiles z-stack images taken from GCaMP6s *Ocm*^*+/+*^ and GCaMP6s *Ocm*^*-/-*^ apical-cochlea at P2. GCaMP6s mice showed endogenous green fluorescence (green). OCM (white) expression can be detected in P2 GCaMP6s *Ocm*^*+/+*^ mice, and exhibited a gradient expression along the cochlear coil, with a higher level at the base and lower at the apex (n = 3). Myo7a (red) was used as the hair cell marker.

### The absence of OCM changes Ca^2+^ signaling in the postnatal development of GCaMP6s Ocm^-/-^ mice

In previous studies, we showed that OCM expression in OHCs influences intracellular Ca^2+^ signaling during postnatal development. By loading dissected cochlear preparations with the Ca^2+^ indicator Fluo-4, we demonstrated that the onset of OCM expression leads to slower KCl-induced Ca^2+^ transients than those elicited in OHCs from *Ocm*^-/-^ littermates (Murtha et al., 2022). Changes in intracellular Ca^2+^ induced by extracellular KCl caused increases in GCaMP6s fluorescence in OHCs (**Figure 3A, Movie 1**). The change in Ca^2+^ levels in OHCs from all 3 rows was probed by the fractional change in signal fluorescence (∆F/F_0_, **Figure 3B**). OHCs from GCaMP6s *Ocm*^*-/-*^ mice showed significantly increased average maximum ∆F/F_0_ compared to GCaMP6s *Ocm*^*+/+*^ OHCs (*P* < 0.001, *t*-test, **Figure 3C**). The time course of the Ca^2+^ transient induced by extracellular KCl in GCaMP6s *Ocm*^*-/-*^ OHCs exhibited a faster rise-time constant compared to GCaMP6s *Ocm*^*+/+*^ OHCs at P2 (**Figure 3D**). These data indicate that GCaMP6s *Ocm*^*-/-*^ mice also show changes in Ca^2+^ signaling. As a Ca^2+^ binding protein, GCaMP6s could alter Ca^2+^ buffering in OHCs. However, GCaMP6s was present in OHCs from both control and *Ocm*^*-/-*^ mice, and Ca^2+^ activity in GCaMP6s *Ocm*^*+/+*^ OHCs was similar to that previously measured in non-GCaMP6s transgenic mice (Murtha et al., 2022).

**Figure 3.**
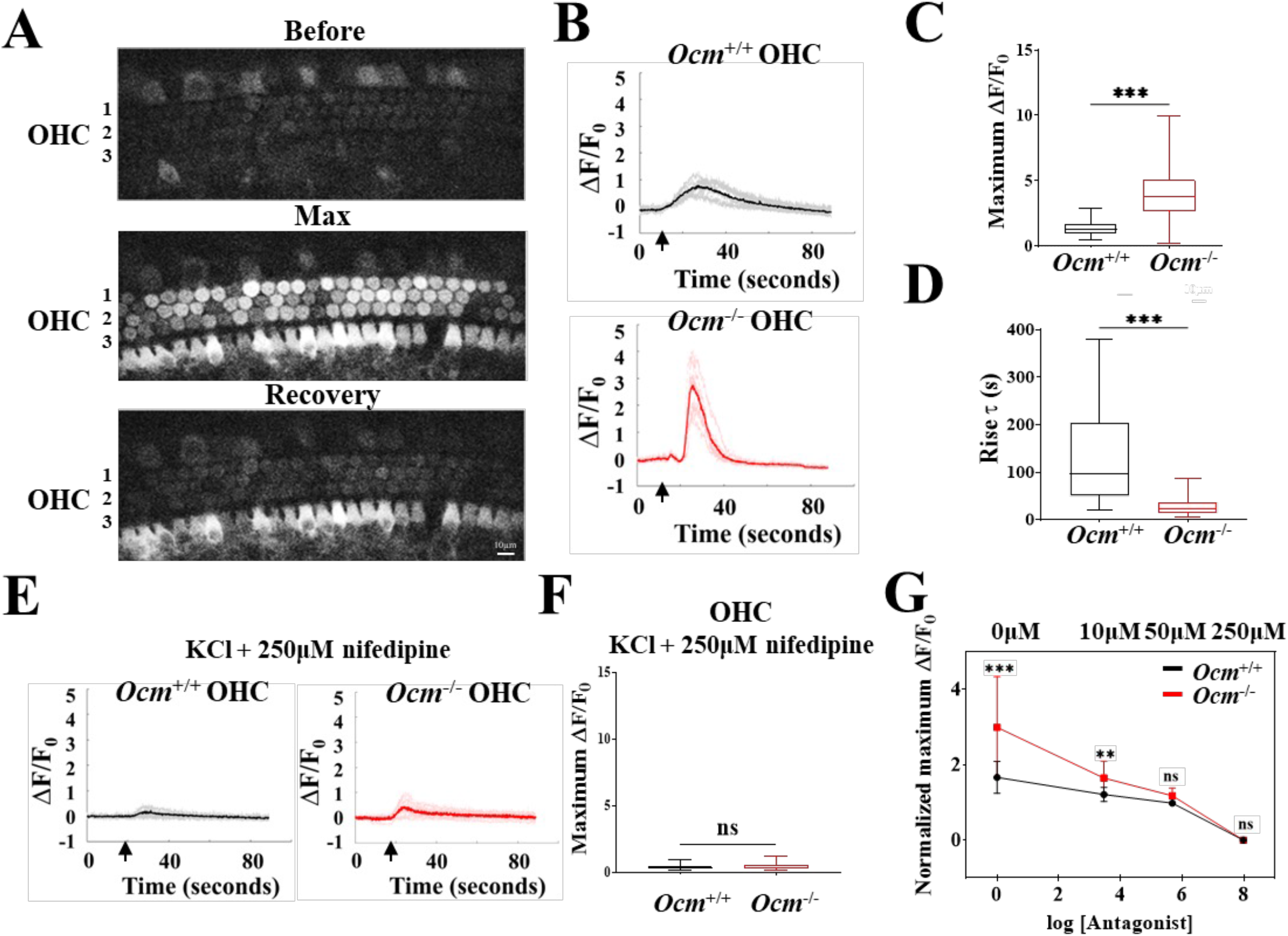
The absence of OCM increased the fractional change of voltage-gated Ca^2+^ influx in OHCs. **A**. Organ of Corti were taken from GCaMP6s mice at P2. KCl (37 mM final concentration) superfusion was administered to elicit Ca^2+^ transients. GCaMP6s fluorescence is shown before the superfusion of KCl (top), at peak response (middle), and the recovery stage (bottom). Recordings were taken at room temperature (∼24℃). **B**. Representative plots of GCaMP6s fluorescence fractional change (ΔF/F_0_) in OHCs induced by KCl superfusion (arrow) for REF s. GCaMP6s *Ocm*^*+/+*^ and GCaMP6s *Ocm*^*-/-*^ mice at P2 were studied. Results show individual ROI fluorescence ∆F/F_0_ traces view (grey and light red) and mean ∆F/F_0_ for OHCs (black and red) from P2 GCaMP6s *Ocm*^*+/+*^ and GCaMP6s *Ocm*^*-/-*^ mice **C**. Average maximum fluorescence intensities from GCaMP6s *Ocm*^*+/+*^ and GCaMP6s *Ocm*^*-/-*^ OHCs at P2 during the application of KCl. Mean peak ΔF/F_0_ are plotted from single OHCs in all 3 rows, n > 3 for each genotype. *** indicates *P* < 0.001, *t*-test. **D**. Rise τ from P2 GCaMP6s *Ocm*^*+/+*^ and *Ocm*^*-/-*^ OHCs during the application of KCl were analyzed, n > 3 for each genotype. ***: P < 0.001, Mann-Whitney test. **E**. Individual ΔF/F_0_ traces (grey and light red) and Mean ∆F/F_0_ (black and red) from GCaMP6s *Ocm*^*+/+*^ and GCaMP6s *Ocm*^*-/-*^ OHCs induced by the KCl application (arrow) with the presence of 250 μM nifedipine. **F**. The mean (± SEM) peak ΔF/F_0_ from GCaMP6s *Ocm*^*+/+*^ and GCaMP6s *Ocm*^*-/-*^ OHCs induced by KCl together with 250 μM nifedipine at P2 was not significant difference (*P* = 0.153, *t*-test). **G**. Dose-response plot of nifedipine antagonism of KCl-induced Ca^2+^ transients in GCaMP6s *Ocm*^*+/+*^ and GCaMP6s *Ocm*^*-/-*^ OHCs at P2. Cochlear explants were incubated with 0, 10 μM, 50 μM and 250 μM nifedipine before KCl perfusion. Relative mean (± SEM) peak ΔF/F_0_ from different nifedipine dosage treatments were normalized to average peak ΔF/F_0_ when 250 μM nifedipine was applied. X-axis: logarithm of nifedipine concentration. *: *P* < 0.05, ***: *P* < 0.001, *one-way ANOVA* followed by Bonferroni’s test.

We then sought to verify whether the KCl-induced Ca^2+^ transients depend on voltage-gated Ca^2+^ channels. In both GCaMP6s *Ocm*^*+/+*^ and GCaMP6s *Ocm*^*-/-*^ OHCs, Ca^2+^ transients were nearly eliminated when KCl was applied together with 250 μM nifedipine (**Figure 3E**), and average maximum ∆F/F_0_ was comparable in OHCs from both genotypes (*P* = 0.153, *t*-test. **Figure 3F**). A normalized dose-response curve (normalized to ∆F/F_0_ with 250 μM nifedipine) showed that KCl-induced Ca^2+^ transients were partially blocked by 10 or 50 μM nifedipine (*P* < 0.001, *t*-test, **Figure 3F**). The dose-dependent response to nifedipine inhibition revealed that voltage-gated Ca^2+^ channels are required for Ca^2+^ transients induced by KCl.

The Ca_v_1.3 channel is the main voltage-gated L-type Ca^2+^ channel expressed in hair cells (Michna et al., 2003; Platzer et al., 2000). We investigated whether the lack of OCM altered the expression of Ca_v_1.3. We performed qRT-PCR, western blots, and immunofluorescence on cochleae from P2 GCaMP6s *Ocm*^*+/+*^ and GCaMP6s *Ocm*^*-/-*^mice. The relative Ca_v_1.3 mRNA expression (encoded by *CACNA1D*) in the GCaMP6s *Ocm*^*-/-*^ cochlea was significantly downregulated compared to GCaMP6s *Ocm*^*+/+*^ cochlea (*P* < 0.001, *t*-test, **Figure 4A**). Western blot also showed that the GCaMP6s *Ocm*^*-/-*^ cochlea exhibited a significantly lower level of Ca_v_1.3 protein compared to GCaMP6s *Ocm*^*+/+*^ cochlea (*P* < 0.05. *t*-test, **Figure 4B-C**).

**Figure 4.**
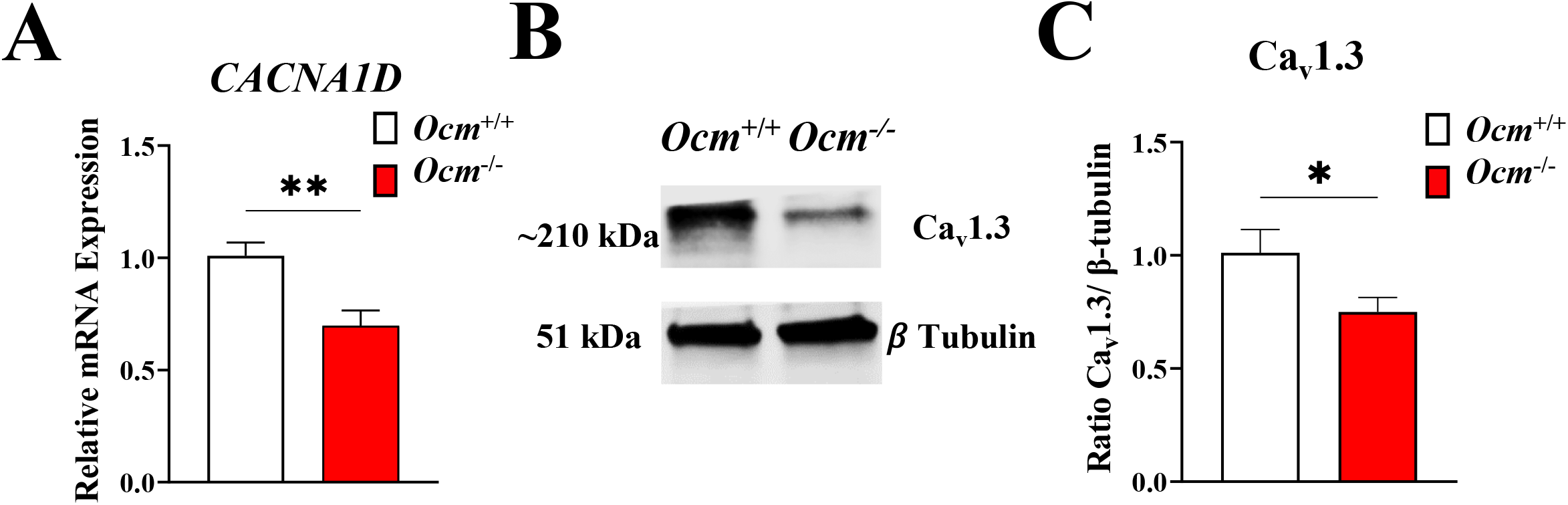
The absence of OCM decreased the expression of Ca_v_1.3 channel in OHCs. **A**. qRT-PCR results of Ca_v_1.3 (*CACNA1D*), from GCaMP6s *Ocm*^*+/+*^ and GCaMP6s *Ocm -*_*/-*_ mice at P2. Total RNA was extracted from GCaMP6s mice cochleae at P2 (n > 3 for each genotype). Plots are the mean (± SEM) normalized mRNA levels relative to GCaMP6s *Ocm*^*+/+*^. Asterisks (*) indicate a significant difference between *Ocm*^*+/+*^ and *Ocm*^*-/-*^ gene expression for the same age. ***: *P* < 0.001, *t*-test. **B, C**. Representative western blot for Ca_v_1.3 protein expression levels detected in cochlea harvested from P2 GCaMP6s *Ocm*^*+/+*^ and GCaMP6s *Ocm*^*-/-*^ mice. Each genotype has n > 3 independent repeats, each replicate contained 3 cochleae and total animal n = 9 for each genotype. *: *P* < 0.05, *t*-test. β-tubulin (loading control) was used for normalization. The plot is the mean (± SEM) normalized grey values relative to GCaMP6s *Ocm*^*+/+*^.*: *P* < 0.05, *t*-test.

### Targeted deletion of Ocm increased spontaneous calcium signaling in the immature cochlea

During early postnatal development, OHCs show spontaneous Ca^2+^ activity that can be synchronized by ATP-induced Ca^2+^ waves originating from the GER, and depends on extracellular Ca^2+^ via Ca_v_1.3 channel (Ceriani et al., 2019; Jeng et al., 2020). Since we recently showed that OCM expression alters intracellular Ca^2+^ signaling in OHCs (Murtha et al., 2022), we hypothesized that the absence of OCM in *Ocm*^*-/-*^ mice could affect the spontaneous Ca^2+^ activity in the OHCs. Apical OHCs from both GCaMP6s *Ocm*^*+/+*^ and GCaMP6s *Ocm*^*-/-*^ P2 mice showed increased coordination of Ca^2+^ activity during the occurrence of a Ca^2+^ wave in the GER (time window 2, yellow), compared to when there were no Ca^2+^ waves (time window 1, green, **Figure 5A-F, Movie 2**). Spontaneous Ca^2+^ activity was nearly eliminated in a Ca^2+^-free medium (**Figure S1A**), and in the presence of 250 µM nifedipine (**Figure S1B**), indicating that spontaneous Ca^2+^ activity in OHCs depends on extracellular Ca^2+^.

**Figure 5.**
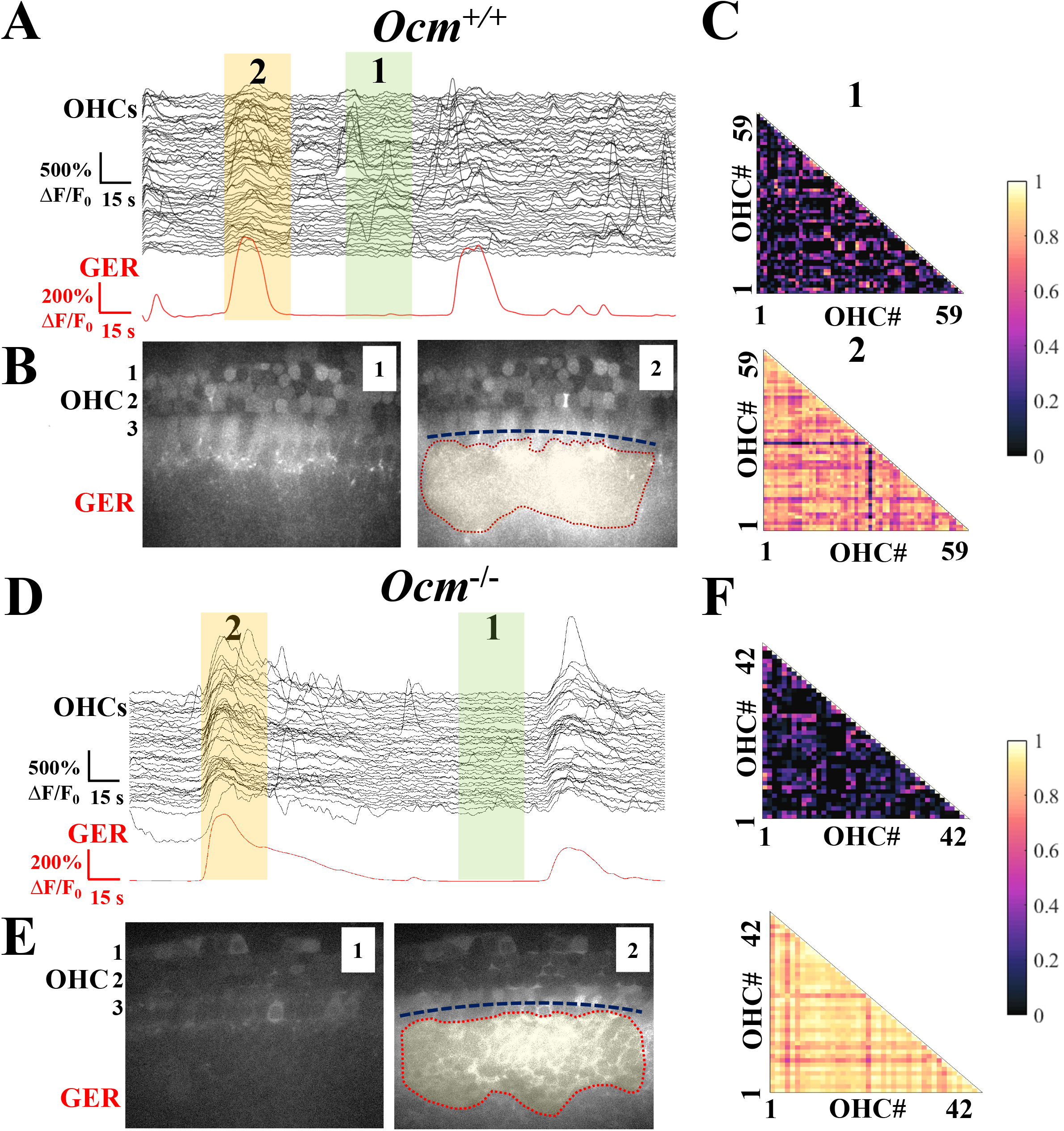
Ca^2+^ waves initiated from the GER synchronized spontaneous Ca^2+^ activities in *Ocm*^*+/+*^ and *Ocm*^*-/-*^ OHCs and DCs. **A, D**. Organ of Corti apical cochlea were taken from GCaMP6s mice at P2. Results show individual ROI GCaMP6s fluorescence intensity fractional change (∆F/F_0_) traces for all OHCs from a single field of view (black), and calcium activity in GER (red). Highlighted green time window (1) represents no Ca^2+^ wave in GER. In contrast, the yellow time window (2) marks the occurrence of a Ca^2+^ wave in GER. Recordings were taken at room temperature (∼24℃). **B, E**. Representative images taken from time windows 1 (green) and 2 (yellow) showing the background and the occurrence of the Ca^2+^ waves in GER for GCaMP6s *Ocm*^*+/+*^ and GCaMP6s *Ocm*^*-/-*^ cochlea, respectively. Dash lines represent the extension size of the Ca^2+^ wave in GER along the cochlear spiral. **C, F**. Representative correlation matrices of ∆F/F_0_ traces in OHCs. Correlation matrices were computed during time window 1 (top panel), and time window 2 (bottom panel). Each matrix element represents Spearman’s rank correlation coefficient (r_s_, see Materials and Methods) of one pair of OHCs from the same cochlear spiral.

To quantify the synchronization of Ca^2+^ signals in OHCs with Ca^2+^ waves in the GER, the average pairwise correlation coefficient between OHC Ca^2+^ traces was calculated in the time window during the occurrence of Ca^2+^ waves in the GER (r_avg_). For OHCs, the r_avg_ from both GCaMP6s *Ocm*^*+/+*^ and GCaMP6s *Ocm*^*-/-*^ showed a positive relationship with the longitudinal extension size of Ca^2+^ activity in the GER and was significantly different from zero (**Figure 6A**, *P* < 0.05 for *Ocm*^*+/+*^, *P* < 0.0001 for *Ocm*^*-/-*^, *f*-test). There was no significant difference between the average extension sizes of Ca^2+^ waves in the GER from either GCaMP6s *Ocm*^*+/+*^ or GCaMP6s *Ocm*^*-/-*^ mice (*Ocm*^*+/+*^: 85.50 ± 61.00 μm, *Ocm*^*-/-*^:

**Figure 6.**
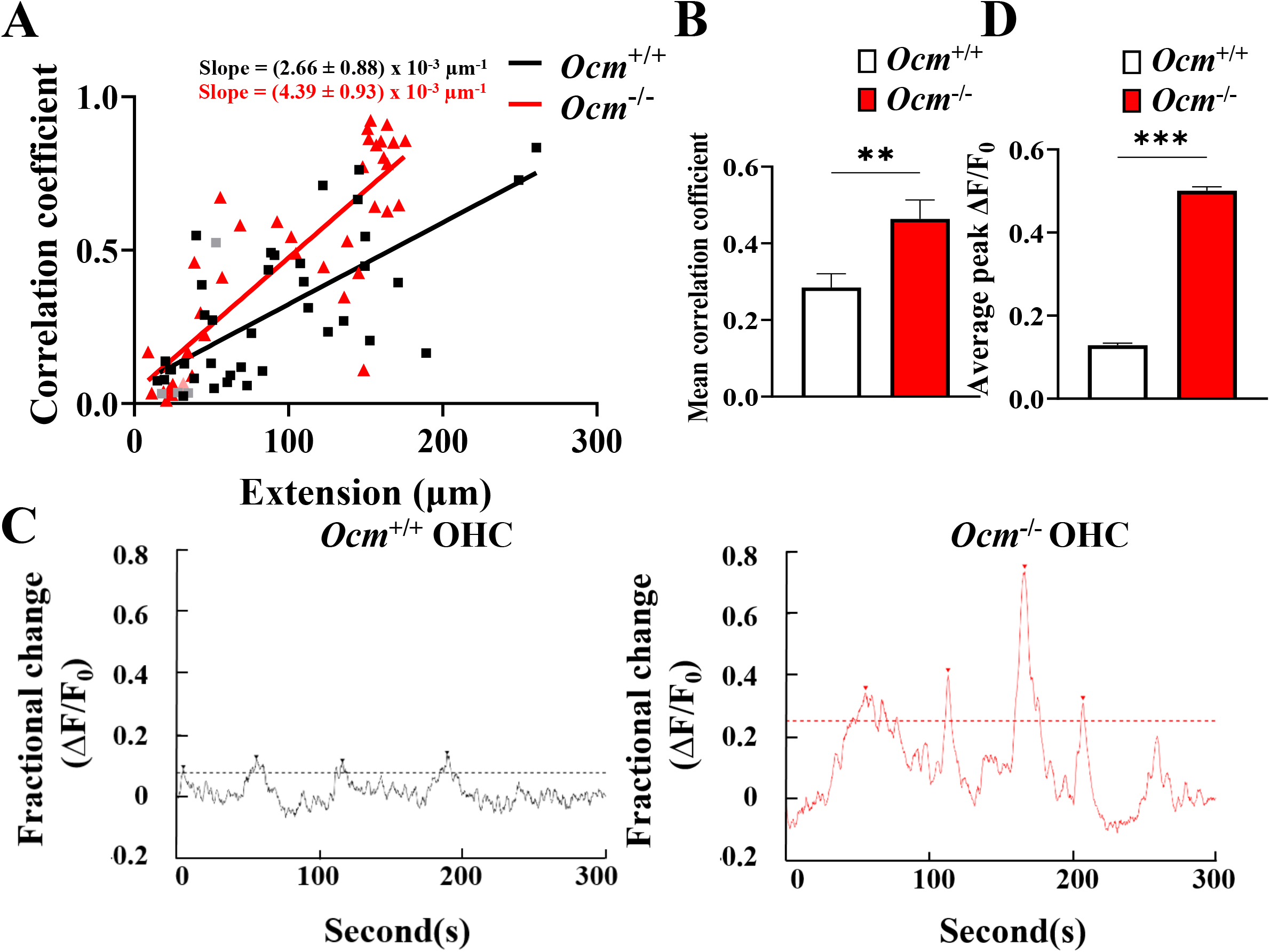
GCaMP6s *Ocm*^*-/-*^ OHCs exhibited a higher level of correlated Ca^2+^ activity and increased maximum ∆F/F_0_ during the Ca^2+^ waves initiated in the GER. **A**. The linear regression between the longitudinal extension of spontaneous Ca^2+^ waves in GER and the average Spearman’s rank correlation coefficient (r_avg_, see Materials and Methods) of OHCs from GCaMP6s *Ocm*^*+/+*^ and *Ocm*^*-/-*^ apical cochlea. Black and red dots symbols represent a significant increase in pairwise OHCs correlation (*P* < 0.05, Mann-Whitney *U*-test) compared to their background time window, while the grey and light red symbols represent the correlation did not increase significantly. The slope rates were shown and were significantly different from zero (*P* < 0.0001, *f*-test).. 11 cochleae, 8 mice for *Ocm*^*+/+*^. 11 cochleae, 6 mice for *Ocm*^*-/-*^ waves per sample, 44 waves from *Ocm*^*+/+*^ cochlea, 42 waves from *Ocm*^*-/-*^ cochlea were calculated. **B**. Average correlation coefficient in between the extension size of Ca^2+^ wave in GER and the r_avg_ in OHCs during the Ca^2+^ waves calculated from above. Mean ± SEM. ns: no significance; **: *P* < 0.01, *Mann-Whitney* test. **C**. Representative Ca^2+^ signaling in single OHC from GCaMP6s *Ocm*^*+/+*^ and GCaMP6s *Ocm*^*-/-*^ apical cochlea, the arrow represents a Ca^2+^ spike in OHC, only spikes that exceeded the threshold (dash line) were calculated (see Materials and Methods). **D**. Average maximum ∆F/F_0_ of OHCs spontaneous Ca^2+^ activity. Mean ± SEM. ***: *P* < 0.001, *Mann-Whitney* test. 2545 Ca^2+^ spikes in OHCs from 11 GCaMP6s *Ocm*^*+/+*^ cochleae and 2303 Ca^2+^ spikes in OHCs from 11 GCaMP6s *Ocm*^*-/-*^ cochleae were calculated.

95.90 ± 60.70 μm, *P* = 0.23, *t*-test). These results indicate that the lack of OCM in OHCs did not change the positive relationship between the extension size of Ca^2+^ waves in the GER and the correlation coefficient (Ceriani et al., 2019). However, GCaMP6s *Ocm*^*-/-*^ OHCs showed a linear regression with a steeper slope (OHCs: 4.39 ± 0.93 nm^-1^) compared to GCaMP6s *Ocm*^*+/+*^ OHCs (2.66 ± 0.88 nm^-1^, *P* < 0.01, *t*-test, **Figure 6A**). We calculated the mean r_avg_ in OHCs. GCaMP6s *Ocm*^*-/-*^ OHCs exhibited a significantly increased mean r_avg_ compared to GCaMP6s *Ocm*^*+/+*^ OHCs (*P* < 0.01, *Mann-Whitney* test, **Figure 6B**).

We then calculated the average maximum ∆F/F_0_ for each Ca^2+^ spike in OHCs (**Figure 6C-D)**. During spontaneous Ca^2+^ waves, the average maximum ∆F/F_0_ in GCaMP6s *Ocm*^*-/-*^ OHCs was greater than in GCaMP6s *Ocm*^*+/+*^ OHCs (**Figure 6D**, *P* < 0.001, *Mann-Whitney* test). Altogether, we found that similar-sized Ca^2+^ waves from the GER produced increased synchronization and higher average maximum ∆F/F_0_ of spontaneous Ca^2+^ activity in the OHCs of GCaMP6s *Ocm*^*-/-*^ relative to those in GCaMP6s *Ocm*^*+/+*^ mice.

### Lack of OCM expression increases ATP-induced Ca^2+^ signaling and purinergic receptors expression in cochlear OHCs

ATP signaling plays a central role during the development of the cochlea. Cochlear cells exhibit a diverse array of purinergic signaling components including all subtypes of ionotropic P2X and metabotropic P2Y receptor subunits (Housley et al., 2009). In the immature cochlea, OHCs express P2X and P2Y receptors and exhibit depolarizing, ATP-gated currents (Bobbin, 2001; Glowatzki et al., 1997).

Initially, we investigated whether purinergic signaling is altered in the absence of OCM, by investigating whether spontaneous Ca^2+^ activity in OHCs was blocked by the ionotropic P2X purinergic receptor antagonist PPADS (100 μM). Both GCaMP6s *Ocm*^*+/+*^ and GCaMP6s *Ocm*^*-/-*^ OHCs showed spontaneous Ca^2+^ activity (**Figure 7A, C**). However, in the presence of PPADS, the Ca^2+^ waves originating from the GER failed to synchronize the Ca^2+^ signaling in OHCs (*P* > 0.05, *Mann-Whitney U*-test, **Figure 7B, D**). The presence of PPADS also affected the linear relationship between the extension size of Ca^2+^ waves in GER and the r_avg_ in both GCaMP6s *Ocm*^*+/+*^ and GCaMP6s *Ocm*^*-/-*^ OHCs (**Figure 7E**, *P* = 0.69 for *Ocm*^*+/+*^ OHCs and *P* = 0.70 for *Ocm*^*-/-*^ OHCs compared to zero, *f*-test). The average maximum ∆F/F_0_ was significantly decreased in both GCaMP6s *Ocm*^*+/+*^ and GCaMP6s *Ocm*^*-/-*^ OHCs (**Figure 7F, G**, *P* < 0.001 for *Ocm*^*+/+*^, *P* < 0.05 for *Ocm*^*-/-*^, *Mann-Whitney* test). We noticed that the average maximum ∆F/F_0_ decreased more in *Ocm*^*-/-*^ OHCs compared to *Ocm*^*+/+*^ OHCs (*Ocm*^*+/+*^: 3.80%, *Ocm*^*-/-*^: 10.53%, *P* < 0.001, *t*-test after normalization).

**Figure 7.**
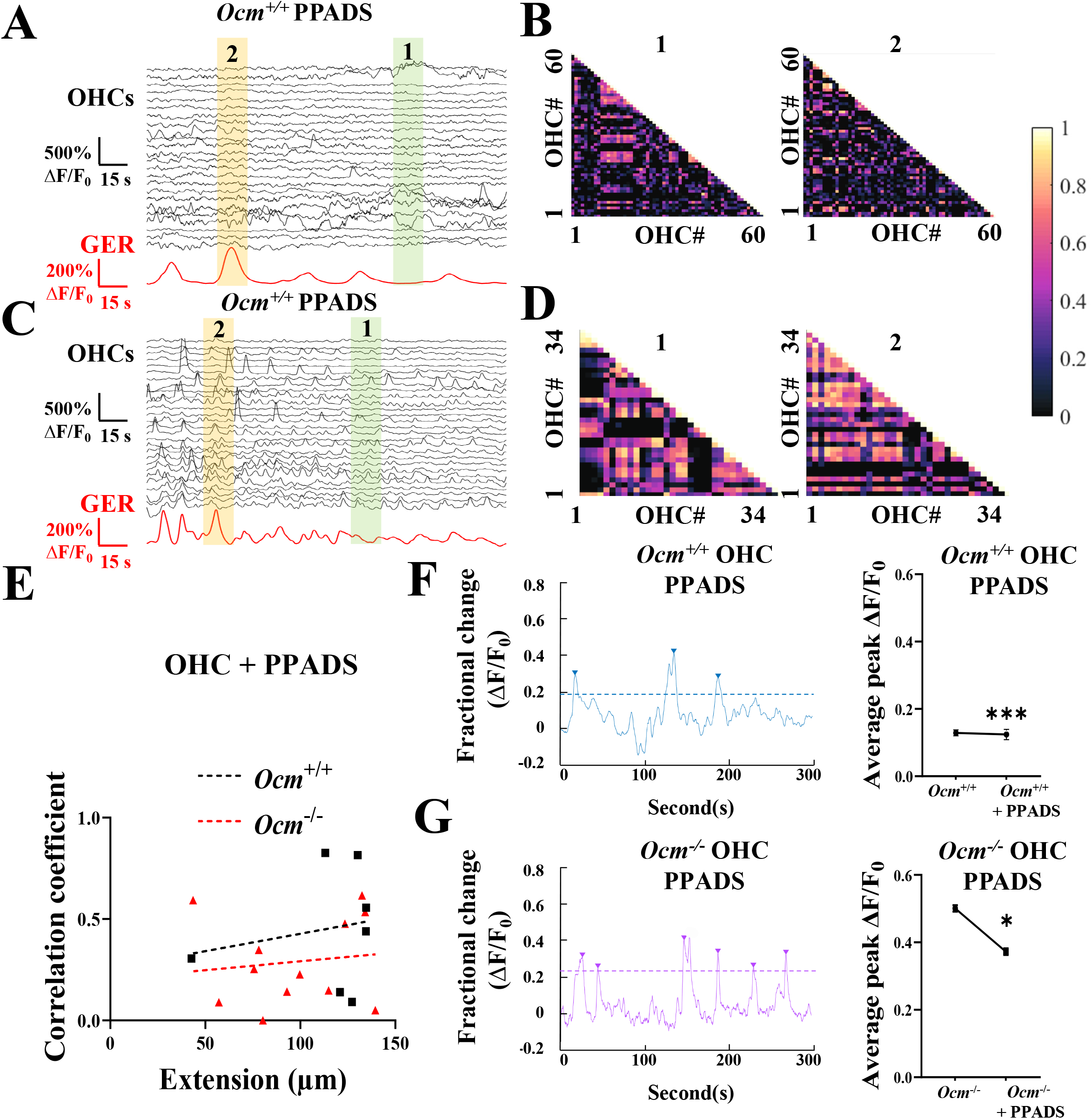
The purinergic receptor is required for the synchronization of spontaneous Ca^2+^activity. **A-D** Individual ROI ∆F/F_0_ traces for all OHCs from a single field of view (black) and calcium activity in GER (red) were taken from GCaMP6s *Ocm*^*+/+*^ and GCaMP6s *Ocm*^*-/-*^ mice at P2. Highlighted green (1) and yellow (2) time windows represent no Ca^2+^ wave (background), and the occurrence of Ca^2+^ wave in GER, respectively, and were used for correlation analysis. The right panel shows representative Ca^2+^ signaling in a single OHC. **B** and **D** show representative correlation matrices calculated from **A** and **C** time window 1 (background, left panel) and time window 2 (during the occurrence of Ca^2+^ wave in GER, right panel). Each matrix element represents Spearman’s rank correlation coefficient (r_s_) of one pair of OHCs. **E** The linear regression between the longitudinal extension of spontaneous Ca^2+^ waves in GER and the average Spearman’s rank correlation coefficient (r_avg_, see Materials and Methods) of OHCs from GCaMP6s *Ocm*^*+/+*^ and GCaMP6s *Ocm*^*-/-*^ apical cochlea with the continuous presence of PPADS (100 µM). The slope from both *Ocm*^*+/+*^ and GCaMP6s *Ocm*^*-/-*^ OHC showed no significant deviation from zero (*P* = 0.68 for *Ocm*^*+/+*^, *P* = 0.69 for *Ocm*^*-/-*^, *f*-test). **F, G**. Representative Ca^2+^ signaling in single OHC from GCaMP6s *Ocm*^*+/+*^ and GCaMP6s *Ocm*^*-/-*^ apical cochlea with the presence of PPADS (100µM). Arrow represents a Ca^2+^ spike in OHC, only spikes that exceeded the threshold (dash line) were calculated (see Materials and Methods). The average maximum ∆F/F_0_ of OHCs spontaneous Ca^2+^ activity with the presence or absence of PPADS was plotted. Mean ± SEM. *: P < 0.05, ***: P < 0.001, animal n = 3, n > 600 OHCs for each genotype were tested, *Mann-Whitney* test.

We then investigated whether Ca^2+^ transients in OHCs elicited by extracellular ATP were affected in the absence of OCM at P2. GCaMP6s *Ocm*^*-/-*^ OHCs showed significantly higher maximum ∆F/F_0_ signal compared to GCaMP6s *Ocm*^*+/+*^ mice (**Figure 8A-C**, *P* < 0.001 for OHCs, *t*-test). ATP-induced Ca^2+^ transients in GCaMP6s *Ocm*^*-/-*^ OHCs were nearly eliminated in Ca^2+^-free medium (**Figure 8D**). The Ca^2+^ activity was abolished in OHCs when ATP was applied with the presence of 100 μM PPADS (**Figure 8E**). Altogether, these data indicate that purinergic signaling can modulate spontaneous Ca^2+^ activity in both GCaMP6s *Ocm*^*+/+*^ and GCaMP6s *Ocm*^*-/-*^ OHCs.

**Figure 8.**
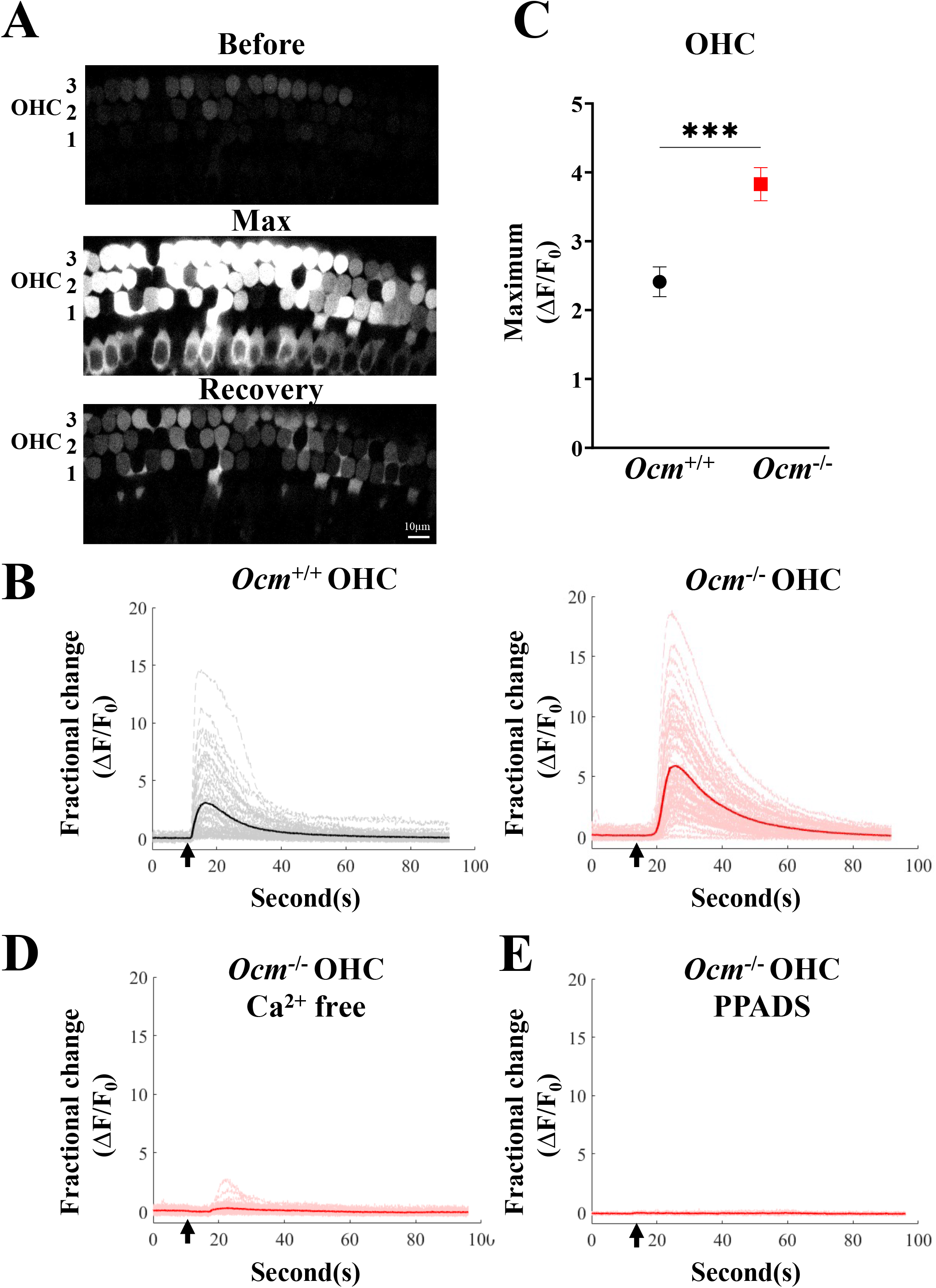
ATP-induced Ca^2+^ transients are altered in *Ocm*^*-/-*^ mice. **A**. Organ of Corti were taken from GCaMP6s mice at P2. Representative results of GCaMP6s fluorescence are shown before 200 μM ATP superfusion, at peak response, and at the recovery stage **B**. Representative plots of the fractional change (ΔF/F_0_) in GCaMP6s fluorescence induced by 200 μM ATP superfusion (arrow). Results show individual ROI fluorescence ∆F/F_0_ traces view (grey and light red) and mean ∆F/F_0_ for OHCs from P2 GCaMP6s *Ocm*^*+/+*^ and GCaMP6s *Ocm*^*-/-*^ mice, n > 3 for each genotype. **C**. Average maximum ΔF/F_0_ of OHCs from GCaMP6s *Ocm*^*+/+*^ and GCaMP6s *Ocm*^*-/-*^ mice at P2, induced by 200 μM ATP superfusion, Animal n > 3. Mean (± SEM). *: *P* < 0.05; ***: *P* < 0.001, t-test. **D, E**. Mean ΔF/F_0_ from GCaMP6s *Ocm*^*-/-*^ P2 OHCs induced by 200 μM ATP (arrow) in Ca^2+^-free medium, or in the presence of the purinergic receptor antagonist, PPADS (100 μM).

Since the lack of OCM causes a higher GCaMP6s fluorescence induced by ATP in OHCs similar to KCl, we investigated whether there were changes in the expression of P2X receptors. Among known P2X receptors, P2X2, P2X3, and P2X7 are all expressed in the cochlea during development (Housley et al., 1998; Huang et al., 2006; Nikolic et al., 2003). We performed qRT-PCR on cochlea harvested from GCaMP6s *Ocm*^*+/+*^ and GCaMP6s *Ocm*^*-/-*^ mice at P2. *P2RX2, P2RX3*, and *P2RX7* mRNA expression were significantly higher in GCaMP6s *Ocm*^*-/-*^ cochlea compared to GCaMP6s *Ocm*^*+/+*^ cochlea (**Figure 9A**, for *P2RX2* and *P2RX7, P* < 0.05, for *P2RX3, P* < 0.001 *t*-test). However, *P2RX2* receptor expression demonstrated the greatest fold change among the three purinergic receptors (>10 fold) in OHCs from GCaMP6s *Ocm*^*-/-*^ mice compared with littermate controls. We then investigated P2X2 protein expression using western blot and immunofluorescence labeling. The western blot results revealed that GCaMP6s *Ocm*^*-/-*^ cochlea showed significantly upregulated P2X2 protein expression relative to GCaMP6s *Ocm*^*+/+*^ (**Figure 9B**, *P* < 0.05, *t*-test). As one of the most abundant purinergic receptors in the inner ear (Kim et al., 2020; Köles et al., 2019), the P2X2 receptor plays an important role in cochlear physiological processes. GCaMP6s *Ocm*^*-/-*^ OHCs exhibited greater P2X2 immunofluorescence labeling compared to GCaMP6s *Ocm*^*+/+*^ (**Figure 9C**).

**Figure 9.**
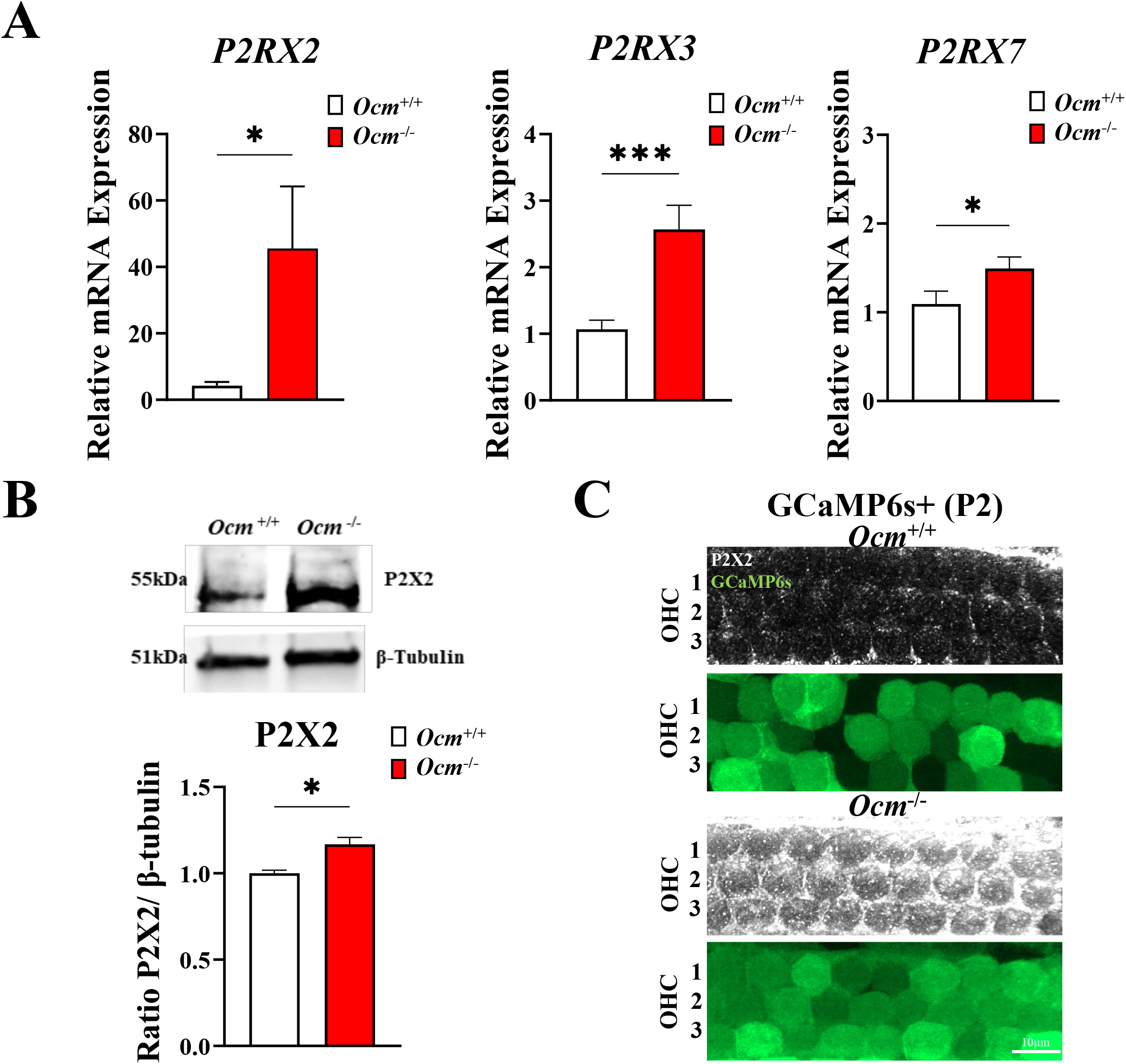
P2X purinoceptor 2 (P2X2) expression is upregulated in *Ocm*^*-/-*^ mice. **A**. qRT-PCR results show that *P2RX2, P2RX3*, and *P2RX7* relative expression level is significantly increased in GCaMP6s *Ocm*^*-/-*^ cochlea at P2 compared to GCaMP6s *Ocm*^*+/+*^. Cochleae were taken from GCaMP6s mice. Whole cochlea spirals were dissected from the cochlea, and total mRNA was extracted immediately after dissection. n > 3 for each genotype. Plot is the mean (± SEM). All values were normalized to *Ocm*^*+/+*^. *: *P* < 0.05, ***: *P* < 0.001, *t*-test. **B**. Representative western blot for P2X2 protein expression levels detected in cochlea derived from P2 GCaMP6s *Ocm*^*+/+*^ and GCaMP6s *Ocm*^*-/-*^ mice. β-tubulin (loading control) was used for normalization. n ≥ 6 for each genotype. Plot is the mean (± SEM) normalized grey values relative to *Ocm*^*+/+*^.* : *P* < 0.05, *t*-test. **C**. Maximum intensity projections of P2X2 immunolabeling on three rows of OHCs harvested from P2 GCaMP6s *Ocm*^*+/+*^ and GCaMP6s *Ocm*^*-/-*^ mice, n ≥ 3 for each genotype. P2X2 (white), and GCaMP6s (green) are shown.

### The number of afferent fibers in Ocm^-/-^ mice is increased

In the cochlea, the type II afferent fibers cross the tunnel and contact multiple OHCs, and form branches to the outer supporting cells, including Deiter’s cells and Hensen’s cells (Fechner et al., 2001). The maturation of synaptic contacts includes a transformation from multiple small to one single presynaptic active zone (Michanski et al., 2019). Thus, we expected that the changes in spontaneous Ca^2+^ activity might affect ribbon synapse maturation and afferent innervation in GCaMP6s *Ocm*^*-/-*^ OHCs. We first counted the number of afferent ribbons from GCaMP6s *Ocm*^*+/+*^ and GCaMP6s *Ocm*^*-/-*^ OHC regions both prior to the onset of hearing and after hearing onset. We performed immunofluorescence at P2, P6, P10, and 3-4 wks in apical cochlear regions using an antibody against C-terminal-binding protein 2 (CtBP2), a marker of the pre-synaptic ribbon. In the OHC region, the number of CtBP2 puncta increased between P2 and P6 in both GCaMP6s *Ocm*^*+/+*^ and GCaMP6s *Ocm*^*-/-*^. Between P6 and P10 there was a drastic reduction of puncta in GCaMP6s *Ocm*^*+/+*^ mice, but not in GCaMP6s *Ocm*^*-/-*^ mice. At P10, GCaMP6s *Ocm*^*-/-*^ apical cochlea showed an increased number of CtBP2 puncta compared to GCaMP6s *Ocm*^*+/+*^ mice (**Figure 10A-E**, *P* < 0.05 *two-way ANOVA*). However, by 3-4 wks, the number of ribbons labeled by CtBP2 in GCaMP6s *Ocm*^*+/+*^ and GCaMP6s *Ocm*^*-/-*^ apical cochlea showed no significant difference. Our results suggested that the synaptic maturation and pruning in GCaMP6s *Ocm*^*-/-*^ cochlea was delayed compared to GCaMP6s *Ocm*^*+/+*^ cochlea.

**Figure 10.**
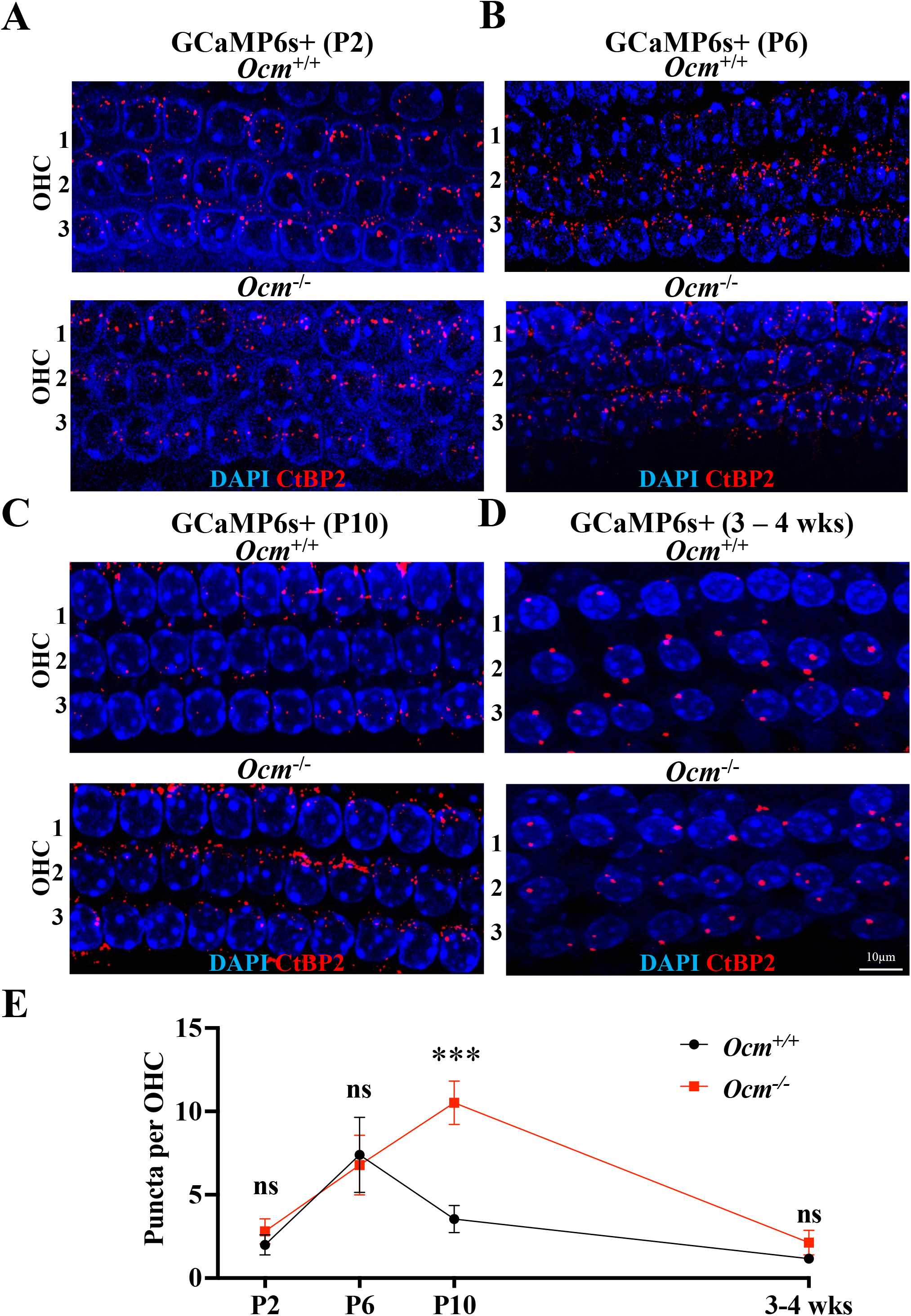
Ribbon synapse maturation is delayed in GCaMP6s *Ocm*^*-/-*^ mice. **A-D**. Maximum intensity projections of confocal z-stacks from GCaMP6s *Ocm*^*+/+*^ and GCaMP6s *Ocm*^*-/-*^ apical cochlea at P2, P6, P10, and 3-4 wks. Three rows of OHCs with ribbon synapses (CtBP2, red) and DAPI (blue) are shown. **E**. The average number of ribbon synapses puncta per OHC from the apical cochlea was measured. The CtBP2 puncta within 4µm of the OHC nuclei were counted. Bar graphs show the mean (± SEM) number of ribbons per OHC. n ≥ 3, ***: P < 0.001, *two-way ANOVA* followed by Bonferroni’s test.

We then used peripherin to examine type II afferent spiral ganglion (SG) fibers (Hafidi, 1998) in cochleae from GCaMP6s *Ocm*^*+/+*^ and GCaMP6s *Ocm*^*-/-*^ mice at P6 and P10 (**Figure 11A, B**). The SG neurons formed outer spiral fibers that terminate on OHCs after long spiral courses. Peripherin immunofluorescence revealed that GCaMP6s *Ocm*^*-/-*^ mice showed a similar number of tunnel crossing fibers compared to GCaMP6s *Ocm*^*+/+*^ mice at P6 (**Figure 11C**, *P* = 0.31, *t*-test), but had an increased number of tunnel crossing fibers compared to GCaMP6s *Ocm*^*+/+*^ mice at P10 (**Figure 11D**, *P* < 0.05, *t*-test). These data provide evidence that the increased spontaneous Ca^2+^ signaling in GCaMP6s *Ocm*^*-/-*^ cochlea changes the maturation of afferent synapses and innervation of afferent fibers during development.

**Figure 11.**
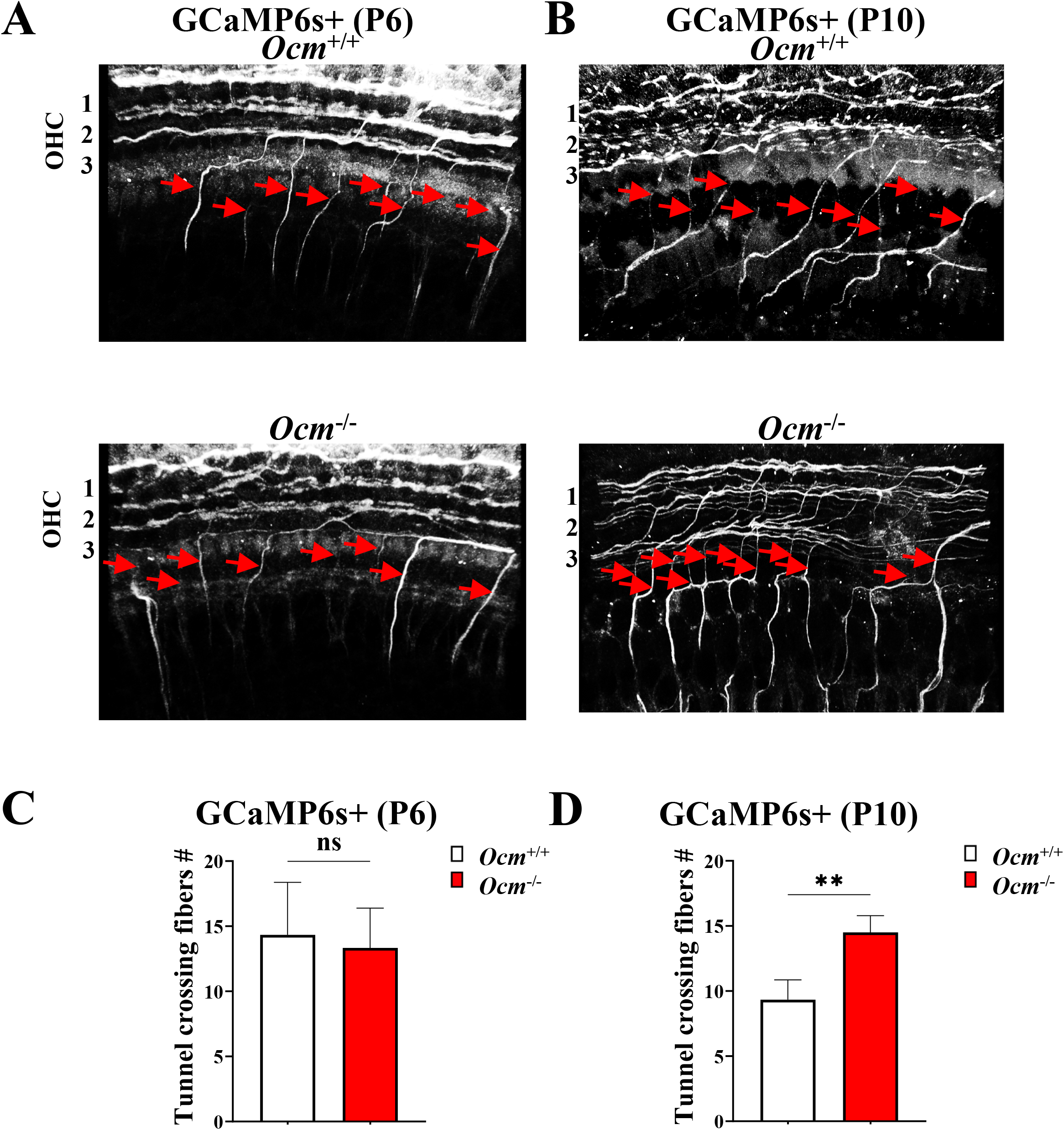
Afferent fibers increased in *Ocm*^-/-^ at P10. **A, B**. Maximum intensity projections of confocal z-stacks from GCaMP6s *Ocm*^+/+^ and GCaMP6s *Ocm*^-/-^ mice at P6 and P10. Afferent fibers labeled with peripherin (white) and DAPI (blue) are shown. The outer spiral fibers of type II spiral ganglion neurons travel toward the cochlear base (arrows). **C, D**. The average number of tunnel crossing afferent fibers from *Ocm*^+/+^ and *Ocm*^-/-^ mice at P6 and P10. Values are means ± SEM, n ≥ 3, **: P < 0.01, t-test.

## Discussion

In the present study, we showed that OCM influences the development of spontaneous activity in OHCs and modulates their neonatal afferent innervation. We generated *Ocm*^*+/+*^ and *Ocm*^*-/-*^ mice with a genetically encoded calcium sensor (GCaMP6s). Similar to other studies (Climer et al., 2021; Tong et al., 2016), GCaMP6s *Ocm*^*-/-*^ mice showed normal hearing at 3-4 weeks of age but exhibited an early onset hearing loss at 7-9 weeks. Based on our previous findings (Murtha et al., 2022), we focused this study at the onset of *Ocm* expression (P2). GCaMP6s *Ocm*^*-/-*^ P2 OHCs have higher maximum ΔF/F_0_ GCaMP6s fluorescence intensity induced by ATP and KCl, which is consistent with higher levels of free cytosolic Ca^2+^ previously reported (Murtha et al., 2022). Both GCaMP6s *Ocm*^*+/+*^ and GCaMP6s *Ocm*^*-/-*^ OHCs exhibited spontaneous Ca^2+^ activity that is synchronized by Ca^2+^ waves initiated from the GER. However, compared to the GCaMP6s *Ocm*^*+/+*^, GCaMP6s *Ocm*^*-/-*^ OHCs had an increased level of coordinated spontaneous Ca^2+^ activity. Further, GCaMP6s *Ocm*^*-/-*^ OHCs exhibited an increased number of presynaptic ribbons and afferent tunnel-crossing fibers just prior to the onset of the hearing. In this study, loss of OCM results in increased spontaneous Ca^2+^ signaling and delayed the maturation of afferent innervation. Taken together OCM contributes to the modulation of Ca^2+^ signaling and the maturation of afferent connectivity in the developing mouse cochlea.

### OCM modulates the expression of Ca^2+^-related genes during development

Purinergic receptors have been implicated in auditory neurotransmission, regulation of cochlear homeostasis, cochlear development, and neurodegenerative conditions (Burnstock, 2016; Housley et al., 2009; Linden et al., 2019). Previous studies have shown that all P2X receptors are transiently expressed in the developing mammalian cochlea (Vlajkovic and Thorne, 2022). Among these purinergic receptors, P2X3 and P2X7 are expressed in sensory hair cells from embryonic day 18 (E18) to P6 (Huang et al., 2006; Nikolic et al., 2003). P2X2 receptors are the predominant purinergic receptor in the mature cochlea and is expressed in hair cells before P15 (Jarlebark et al., 2002). We found that the expression of P2X2, P2X3, and P2X7 purinergic receptors were all upregulated due to the lack of OCM. The upregulation of P2X receptors in the *Ocm*^-/-^ cochlea could be explained either by an OCM-mediated regulatory pathway or by the effect of Ca^2+^ levels on purinergic reveptors expression. Several studies suggest CaBPs may interact directly with purinergic receptors. Roger et al. (2008) found that P2X7 receptors contain a large intracellular C-terminal domain with a Ca^2+^-dependent calmodulin (CaM) binding motif. Sander et al. (2022) reported that Ca^2+^-CaM binding changes the conformation of P2X7, indicating a possible intracellular regulatory pathway of P2X7 receptors. Since OCM and CaM share some functional similarities (Climer et al., 2019; MacManus et al., 1982), it is possible that OCM may interact with P2X receptors to alter their function and expression. Alternatively, upregulated purinergic receptors could be associated with the higher concentrations of cytosolic free Ca^2+^ in *Ocm*^-/-^ OHCs. Loss of OCM or noise exposure can cause increased levels of intracellular free Ca^2+^, leading to Ca^2+^ overloading in OHCs (Murtha et al., 2022; Zuo et al., 2008). Noise exposure also leads to the upregulation of P2X receptors in sensory hair cells (Wang et al., 2003). Thus, the loss of OCM could lead to unrestrained P2X function and increased expression either because of the loss of a direct regulator, or through unfettered cytosolic free Ca^2+^.

Unlike the upregulation of P2X receptors, we found that the expression of Ca_v_1.3, which is the predominant voltage-gated Ca^2+^ channel in hair cells (Hafidi and Dulon, 2004; Michna et al., 2003), was downregulated in the *Ocm*^-/-^ cochlea. Spontaneous Ca^2+^ activity in OHCs is dependent upon the expression of Ca_v_1.3 channels (Ceriani et al., 2019; Jeng et al., 2020). Although there is little evidence for CaBPs directly modulating Ca_v_1.3 expression, previous studies show that CaBPs can modulate Ca_v_1.3 activity. CaM modulates Ca_v_1.3 channel open probability based on cytosolic Ca^2+^ levels (Johny et al., 2013). CaM increases the activity of Ca_v_1.3 channels at low cytosolic Ca^2+^ levels, and decreases the permeability of Ca_v_1.3 channels at high Ca^2+^ levels. Additionally, other CaBPs (e.g., CaBP1, CaBP2, CaBP3, and CaBP4) enhance Ca^2+^ feedback to Ca_v_1.3 channels (Cui et al., 2007). Similar to other CaBPs, OCM could, therefore, modulate the function of Ca_v_1.3 channels. In this way, the expression of OCM or the downregulation of Ca^2+^ entry prevents Ca^2+^ overloading in OHCs and protects OHCs from the deleterious consequences of high Ca^2+^ level. Indeed, mice lacking Ca_v_1.2, another voltage-gated Ca^2+^ channel expressed in hair cells, reduces vulnerability to noise (Zuccotti et al., 2013). The progressive hearing loss phenotype observed in *Ocm*^-/-^ mice could be the result of the insufficiency of protective mechanisms in *Ocm*^-/-^ OHCs to reduce intracellular Ca^2+^ and thus protect mice from hearing loss.

Taken together, our data suggest that OCM regulates the expression of both purinergic receptors and Ca_v_1.3 channels during the early stages of development. Therefore, OCM may contribute significantly to shaping Ca^*2+*^ dynamics, a cornerstone of auditory hair cell function. Since noise leads to changes in Ca^2+^ activity and Ca^2+^-related gene expression, OCM could play a key role in preventing OHC damage due to noise exposure.

### Lack of OCM alters spontaneous Ca^2+^ activity and afferent maturation in OHC

Our data show that *Ocm*^*-/-*^ OHCs exhibited higher synchronized Ca^2+^ activity compared to *Ocm*^*+/+*^ OHCs during development. Spontaneous Ca^2+^ waves are generated from the GER, and these waves travel to the lesser epithelial ridge (LER) where they synchronize Ca^2+^ activity in nearby OHCs via the release of ATP (Ceriani et al., 2019). Interestingly, expression of OCM in OHCs increases during development (Hackney et al., 2005; Simmons et al., 2010), and parellels the downregulation of spontaneous Ca^2+^ activity in developing OHCs (Ceriani et al., 2019; Jeng et al., 2020). In the present study, we found a gradient expression of OCM in OHCs along the tonotopic axis of the cochlea. Indeed, spontaneous Ca^2+^ activity is higher in apical compared to basal OHCs at early postnatal ages (Ceriani et al., 2019; Jeng et al., 2020; Lelli et al., 2009). Taken together, these studies support OCM regulates spontaneous Ca^2+^ activity during the maturation of OHCs.

During the early postanal period, Ca^2+^ influxes and periodic Ca^2+^ stimulation are required for synaptic maturation and afferent refinement (Balland et al., 2006; Sheets et al., 2012; Spitzer, 2006; Tritsch et al., 2007). Recent studies suggest that coordinated Ca^2+^ activity in OHCs is necessary for the formation of their afferent connectivity. The reduction of synchronized spontaneous Ca^2+^ activity in the connexin 30 knockout (*Cx30*^-/-^) OHCs results in a decreased number of ribbon synapses and type II afferent fibers (Ceriani et al., 2019; Jeng et al., 2020). Here, we found that a higher level of spontaneous Ca^2+^ activity at least transiently increases the number of synaptic ribbons and type II afferent fibers. These data reveal that similar to *Cx30*^-/-^, OCM-regulated spontaneous Ca^2+^ activity is also critical to the early patterns of synaptic maturation and afferent innervation during cochlear development.

In summary, the lack of OCM downregulates Ca_v_1.3 channels in OHCs and upregulates P2X2 receptors in the cochlea. Without OCM expression at P2, spontaneous Ca^2+^ activity in OHCs is higher and more synchronized with the GER. We conclude that the lack of OCM changes Ca^2+^ signaling in immature OHCs, resulting in delayed synaptic pruning and changed afferent innervation during the pre-hearing period. We propose that OCM prevents Ca^2+^ overloading and regulates Ca^2+^ signaling necessary for the correct synaptic maturation and afferent innervation during development.

## Materials and Methods

### Animals

Animals were bred at the Baylor University Vivarium. The animal work was licensed by the Institutional Animal Care and Use Committee (IACUC), Baylor University as established by U.S. Public Health Service.

For these studies, *Atoh1*-GCaMP6s mouse lines were maintained on mixed backgrounds. We first crossed *Ocm* wildtype (*Ocm*^*+/+*^) and *Ocm* knockout (*Ocm*^*-/-*^) mice with B6;129S-*Gt(ROSA)26Sor*^*tm96*.*1(CAG-GCaMP6s)Hze*^/J, Ai96(RCL-GCaMP6s) or Ai96, which contains a floxed-STOP cassette preventing transcription of the GCaMP6 slow variant Ca^2+^ indicator. These mice were then crossed with knock-in transgenic mice expressing Cre recombined from the *Atoh1* locus (Chen et al., 2013). *Atoh1*-driven Cre GCaMP6s mice showed tissue-specific expression of endogenous green fluorescence (Cox et al., 2012; Mulvaney and Dabdoub, 2012; Yang et al., 2010). We utilized GCaMP6s positive (GCaMP6s) mice to monitor intracellular Ca^2+^ activities directly after acute dissection.

### Cochlear Function Assays

For measurement of distortion product otoacoustic emissions (DPOAEs), adult mice were anesthetized with xylazine (20 mg/kg, i.p.) and ketamine (100 mg/kg, i.p.). Acoustic stimuli were delivered using a custom acoustic assembly previously described (Maison et al., 2012). Briefly, two electrostatic earphones (EC-1, Tucker Davis Technologies) were used to generate primary tones and a Knowles miniature microphone (EK-3103) was used to record ear-canal sound pressure. Stimuli were generated digitally with 4 s sampling. Ear-canal sound pressure and electrode voltage were amplified and digitally sampled at 20s for analysis of response amplitudes. Both outputs and inputs were processed with a digital I-O board (National Instruments PXI-4461). For measurement of DPOAEs at 2f1 – f2, the primary tones were set so that the frequency ratio, (f2/f1), was 1.2 and so that f2 level was 10 dB below f1 level. For each f2/f1primary pair, levels were swept in 10 dB steps from 20 dB SPL to 80 dB SPL (for f2). At each level, both waveform and spectral averaging were used to increase the signal-to-noise ratio of the recorded ear-canal sound pressure, and the amplitude of the DPOAE at 2f1 – f2 was extracted from the averaged spectra, along with the noise floor at nearby points in the spectrum. Iso-response curves were interpolated from plots of DPOAE amplitude vs. sound level. The threshold was defined as the f2 level required to produce a DPOAE at 0 dB SPL. Right ears were used for all hearing tests.

### Tissue preparation

Cochleae were harvested from *Ocm*^+/+^ or *Ocm*^-/-^ mice of either sex at postnatal day 2 (P2), P10 and 3 - 4 weeks. Pups were euthanized by rapid induction of hypothermia for 2 - 4 min on ice until loss of consciousness. After decapitation, apical coil OHCs were dissected from the organ of Corti in an extracellular medium composed of (in mM): 136.8 NaCl, 5.4 KCl, 0.4 KH_2_PO_4_, 0.3 Na_2_HPO_4_, 0.8 MgSO_4_, 1.3 CaCl_2_, 4.2 NaHCO_3_, 5 HEPES and 5.6 glucose. The pH was adjusted to 7.4 - 7.5 (osmolality ∼306 mmol/kg). The apical coil was then transferred to a small microscope chamber with nylon mesh fixed to a stainless-steel ring on the bottom. The dissected apical coil was immobilized under nylon mesh and visualized using an upright microscope (Leica, DM 6000 FS, Germany) with a 63x / 0.90 water immersion long working distance objective (Leica, 15506362 HCX APO L 63x/0.90 W U-V-I CS2).

### Confocal Ca^2+^ imaging

Ca^2+^ signals from GCaMP6s were recorded using a spinning disk confocal microscope system containing X-Light V2 spinning disk confocal (89 North Inc.) and PRIME95B Photometrics Cooled SMOS with 95% QE (1200 ×1200 pixels with 792 dpi, Teledyne photometric). Images were analyzed offline using ImageJ (NIH). Ca^2+^ signals were measured as relative changes of fluorescence emission intensity (ΔF/F_0_) and calculated by MATLAB. ΔF = F - F_0_, where F is fluorescence at time t and F_0_ is the fluorescence at the onset of the recording.

For spontaneous Ca^2+^ activity, GCaMP6s positive cochlea was recorded immediately after acute dissection at room temperature with 480nm excitation wavelength. Each GCaMP6s fluorescence recording consisted of 1103 frames taken at 6 frames per second from 912 ×912 pixels region, 80 ms exposure time. Fluorescence traces were computed as pixel averages from each OHC and ΔF/F_0_ was calculated in MATLAB. The average GCaMP6s fluorescence of the first 50 frames was used as baseline (F_0_). A Savitzky-Golay filter was then applied to smooth the ΔF/F_0_ (window length = 11, polynomial order = 1). A spike inference algorithm in MATLAB was used to estimate the spike count and position. Only the peak above 4 standard deviations of baseline was caculated (F_0_ + 4 × SD). The minimum peak distance was 5 frames. The average frequency was then computed by dividing the number of spikes by the total duration of the recording. For the spontaneous activity in the GER, the ROI was drawn around the maximum extension of each multicellular calcium event. Fluorescence traces were computed as pixel averages. The longitudinal extension in GER along the cochlear spirals was measured using ImageJ, a stage micrometer calibration slide was used (Olympus, Japan). We calculated the pairwise Spearman’s rank correlation coefficient (r_s_) between every pair of OHCs in the field of view, and pairwise OHCs from the same trial. The significance was tested using Mann-Whitney U-test (one-sided) after Fisher’s z-transformation.

KCl depolarization solution contained (in mM): 142.4 KCl, 0.4 KH_2_PO_4_, 0.3 Na_2_HPO_4_, 0.8 MgSO_4_, 1.3 CaCl_2_, 4.2 NaHCO_3_, 5 HEPES and 5.6 glucose. The pH was adjusted to 7.4 - 7.5 (osmolality ∼305 mmol/kg). ATP depolarization solution was diluted from 100 mM stock ATP (Thermo Fisher Scientific, USA) with the extracellular solution (800 µM ATP for working solution). PPADS (Sigma, USA) and nifedipine (Sigma, USA) were prediluted with DMSO and then diluted using the extracellular solution. Depolarization solution was delivered to the chamber using a PMD-D35 perfusion/media exchange system (Tokai Hit, Japan), followed by 2 min perfusion with the extracellular solution.

For Ca^2+^ transients, 100 µl of KCl depolarization solution, or 100 µl of ATP depolarization solution was added into a microscope chamber containing 300 µl extracellular solution (1:4 dilution, 37 µM KCl final concentration, or 200 µM ATP final concentration in the chamber). The perfusion system (Tokai Hit, Japan) started 8 s after imaging started without generating any turbulence. We measured Ca^2+^ fluorescence from activated OHCs (reach maximum fluorescence intensity between 8-60 s) from all trials separately to avoid the disturbance from spontaneous Ca^2+^ activity.

Each Ca^2+^ fluorescence recording includes 1500 frames taken at 80 frames per second using VisiView (VISITRON, USA). After background subtraction, the activated OHCs were computed as pixel averages using ImageJ. Calcium transient time constant (τ) was calculated using nonlinear regression in MATLAB following these equations:

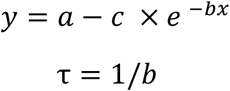

x is the independent variable (time) while y represents fluorescence intensity from each time point.

### Genotyping and qRT-PCR

DNA was extracted from mice tail samples using Extract-N-Amp™ Tissue PCR Kit (Sigma, USA). PCR primers used for genotyping are listed below: *Atoh1*-Cre primer pair forward: 5’-CCGGCAGAGTTTACAGAAGC-3’, reverse: 5’-ATG TTT AGC TGG CCC AAA TG-3’; Cre control primer pair forward: 5’-CTA GGC CAC AGA ATT GAA AGA TCT-3’; reverse: 5’-GTA GGT GGA AAT TCT AGC ATC ATC C-3’; GCaMP6s primer pair forward: 5’-ACG AGT CGG ATC TCC CTT TG - 3’; reverse: 5’- AGA CTG CCT TGG GAA AAG CG - 3’; *Ocm* primer pair forward: 5’- CTC CAC ACT TCA CCA AGC AG - 3’, reverse: 5’- TTT CAT GTT CAG GGA TCA AGT G - 3’; *Ocm* deletion primer pair forward: 5’- CTC CAC ACT TCA CCA AGC AG - 3’, reverse: 5’- GCT TGG GGA CCC CCT GTC TTC A - 3’.

Cochlea from bilateral cochlea were acutely dissected after anesthesia and transferred to the lysis buffer. Total RNA was extracted using RNeasy plus Micro kits (QIAGEN, USA). iScript™ Advanced cDNA Synthesis Kit (BIO-RAD, USA) was used for reverse transcription. qRT-PCR was performed using the SYBR Green PCR Master Mix Kit (BIO-RAD, USA). The *b2m* gene was used as a reference gene (Melgar-Rojas et al., 2015). Quantification of expression (fold change) from the Cq data was calculated following the ΔΔCq method (Schmittgen and Livak, 2008), and normalized to the Cq value in *Ocm*^+/+^ at P0. Briefly, the expression level of a target gene was first normalized to the average level of the corresponding reference gene to obtain the ΔCq value. Then, the ΔΔCq of each gene was calculated as follows: ΔCq (target gene) - ΔCq (target gene from P0 *Ocm*^+/+^ group). 2–ΔΔCq was calculated then to represent the relative expression (fold change).

Primers used for qRT-PCR are as follows: *b2m* forward 5’-TGGTCTTTCTGGTGCTTGTC-3’ and reverse 5’-GGG TGG AAC TGT GTT ACG TAG-3’; *Ocm* forward 5’-ATG AGC ATC ACG GAC ATT CTG AGC-3’ and reverse 5’-CTG GCA GAC ATC TTG GAG AGG C-3’; *CACNA1D* forward 5’-GCA AAC TAT GCA AGA GGC ACC AGA C -3’ and reverse 5’-CTT TGG GAG AGA GAT CCT ACA GGT G -3’; *P2RX2* forward 5’-GCG TTC TGG GAC TAC GAG AC -3’ and reverse 5’-ACG TAC CAC ACG AAG TAA AGC -3’ (PrimerBank ID 27544798a1). *P2RX3* forward 5’-CAA CAC AAC AAG TTT GAA CCC AGC -3’ and reverse 5’-AGG CTT CTT TAG CTT CTC ACTG -3’, *P2RX7* forward 5’-CCC TGC ACA GTG AAC GAG TA -3’ and reverse 5’-CGT GGA GAG ATA GGG ACA GC -3’.

### Immunofluorescence microscopy

For histological analysis and immunocytochemistry, cochleae from neonatal mice were flushed with 4% PFA and then rotated in fixative overnight at 4°C. Cochlea were then separated from the modiolus, followed by 3 washes in PBS, then blocked in 5% normal horse serum (NHS) for 1 h at room temperature.

Samples were stained with antibodies to OCM (Santa Cruz sc-7446, 1:200), P2X2 (alomone lab APR-003, 1:400), CtBP2 (BD Biosciences #612044, 1:200) and peripherin (Sigma AB1530, 1:200). Primary antibodies were incubated overnight at 37ºC. Appropriate Alexa Fluor (Thermo 1:200) and Northern Lights (R&D Systems 1:200) conjugated secondary antibodies were incubated for 1 hour at 37ºC. Slides were prepared using Vectashield mounting media with DAPI (Vector Labs). Images were acquired using the LSM800 microscope (Zeiss) using a high-resolution, oil-immersion objective (Plan-Apochromat 63x, 1.4 NA). Cohorts of samples were immunostained at the same time and imaged under the same optical conditions to allow for direct comparison.

### Western Blotting

Dissected cochleae were harvested (3 cochleae in the same tube, total n = 9 for each genotype) and immediately placed into a lysis buffer containing: 1% Triton X-100, 25 mM Tris, pH 7.4, 150 mM NaCl, 1 mm DTT, 1 mm MgCl_2_, 1 mM phenylmethylsulfonyl fluoride, and 1x protease inhibitor cocktail (Thermo Scientific™). Samples were incubated on ice for 10 minutes and vigorously vortexed twice in 15 second intervals. Lysates were spun down at 4°C 12,000 x g for 20 minutes. Total protein concentration was determined using a BCA Protein Assay Kit (Pierce™, ThermoFisher Scientific, Waltham, Massachusetts). The supernatant was diluted in sample buffer (BioRad, 355 mM β-ME added) and heated for 30 minutes at 37°C. Samples (25 µg total protein) were subject to SDS-PAGE and protein was transferred to a PVDF membrane. Membranes were blocked with 2.5% fish gelatin for 1 hour at RT and incubated overnight at 4°C with primary antibody to rabbit P2X2R (1:400; Alomone Labs, Jerusalem, Israel), mouse Ca_v_1.3 (1:100; ThermoFisher Scientific, Waltham, Massachusetts), and rabbit β-tubulin (1:1000). After washing, membranes were incubated with appropriate HRP-conjugated secondaries (anti-rabbit 1:7,500, anti-mouse 1:100) for 1 hour at RT. Clarity Max ECL Substrate (BioRad, Hercules, California) was used to detect chemiluminescence via a ChemiDoc imaging system. Relative protein expression levels (normalized to β-tubulin) were determined by densitometric analysis using FIJI software.

### Statistical analysis

Statistical analysis was performed using GraphPad Prism 9 and MATLAB software. Statistical comparisons of means were made by t-test or, when normal distribution could not be assumed, the Mann-Whitney *U*-test. For multiple comparisons, one-way or two-way ANOVA followed by Bonferroni’s test was used. Animals of either sex were randomly assigned to the different experimental groups. No statistical methods were used to define the sample size, which was selected based on previous published similar work from our laboratories. Animals were taken from multiple cages and breeding pairs.

**Movie 1**. Representative movie shows KCl-induced Ca^2+^ transient in OHCs.

**Movie 2**. Representative movie shows spontaneous Ca^2+^ activity in GCaMP6s cochlea after acute preparation.

**Figure S1. Extracellular Ca**^**2+**^ **is required for the spontaneous Ca**^**2+**^ **in OHC**. A, B. Representative ΔF/F0 results show individual ROI ∆F/F0 during spontaneous Ca^2+^ activity for all Ocm-/-OHCs (A) in Ca^2+^-free medium or (B) incubated with voltage-gated Ca^2+^ channel blocker nifedipine (250 μM). Right panels show representative Ca^2+^ signaling in a single OHC, no spikes exceeded the threshold in Ca^2+^ media or with the presence of nifedipine (dash line, see Materials and Methods). Recordings were made at room temperature.

**Movie 3**. Representative movie shows extracellular ATP-induced Ca^2+^ transient in OHCs.

Conceptualization Data curation Formal Analysis Funding acquisition Investigation Methodology Project administration. Resources Software. Supervision Validation Visualization Writing — original draft Writing — review & editing

## Additional information

### CRediT authorship contribution statement

**Yang Yang**: Formal Analysis, Investigation, Methodology, Software, Visualization, Writing – original draft, Writing – review & editing. **Kaitlin E. Murtha**: Investigation, Methodology, Writing – review & editing. **Leslie K. Climer**: Methodology, Validation, Writing – review & editing. **Federico Ceriani**: Methodology, Validation, Formal analysis, Writing – review & editing. **Pierce Thompson**: Investigation, Methodology, Writing – review & editing. **Aubrey J. Hornak**: Project administration, Investigation, Writing – review & editing. **Walter Marcotti**: Conceptualization, Methodology, Software, Resources, Writing – review & editing, Supervision, Project administration, Funding acquisition. **Dwayne D. Simmons**: Conceptualization, Methodology, Formal analysis, Investigation, Resources, Data curation, Writing – original draft, Writing – review & editing, Visualization, Supervision, Project administration, Funding acquisition.

All authors approved the final version of the paper. All authorsagree to be accountable for all aspects of the work in ensuringthat questions related to the accuracy or integrity of any part ofthe work are appropriately investigated and resolved. All personsdesignated as authors qualify for authorship, and all those whoqualify for authorship are listed.

