## Supplementary figures and images for "Oncomodulin Regulates Spontaneous Calcium Signaling and Maturation of Afferent Innervation in Cochlear Outer Hair Cells"

### Supplemental Figure 1

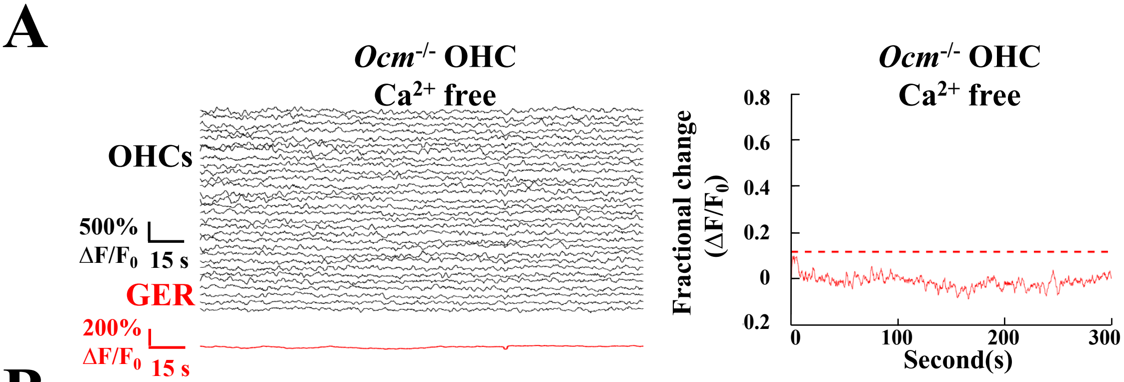

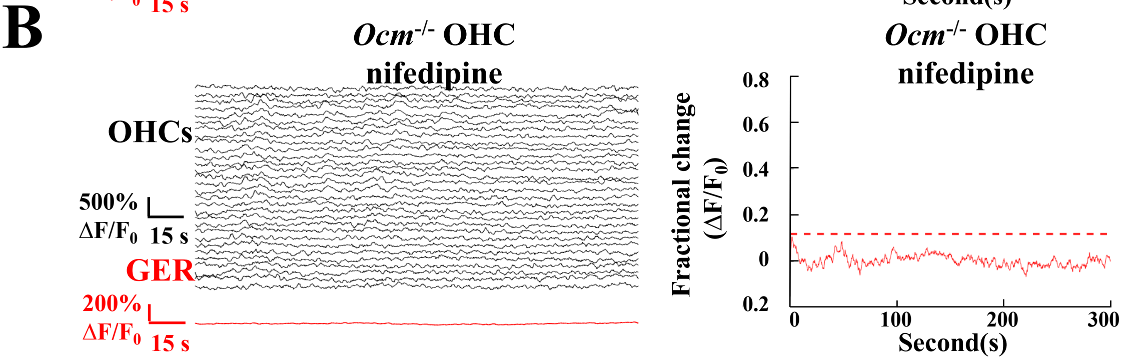
